# Drug-induced versus non-pharmacological wakefulness: similar or different states? A whole brain analysis in TRAP2 transgenic mice

**DOI:** 10.1101/2025.01.29.635430

**Authors:** Renato Maciel, Justin Malcey, Amarine Chancel, Théo Brunel, Patrice Fort, Claudio Marcos Queiroz, Pierre-Hervé Luppi

## Abstract

A large body of data indicate that the aminergic, cholinergic and hypocretin/orexin neurons are responsible for inducing wakefulness. However, recent data showed that other systems might also play a key role. Further, wakefulness induced by different drugs versus non-pharmacological means could be generated by different populations of neurons. To address these questions, we evaluated at the whole brain level in the same mice using TRAP2 model whether the same neurons were activated by the wake-inducing drugs modafinil and solriamfetol versus non-pharmacological wake. Our results show that several subcortical structures such as the bed nucleus of the stria terminalis, central amygdalar nucleus, paraventricular hypothalamic and thalamic and supraoptic nuclei, lateral parabrachial nucleus and lateral reticular area (including its noradrenergic neurons) are significantly more activated by solriamfetol than modafinil and non-pharmacological wakefulness. In contrast, a second category of structures including the orexin neurons, the parasubthalamic and laterodorsal tegmental nucleus are strongly activated in all types of induced wake. Further, some classical wake systems like the dopaminergic neurons of the ventral tegmental area or the dorsal raphe nucleus and the noradrenergic neurons of the locus coeruleus are either very poorly or not strongly activated. These results reveal that many structures not previously involved in wakefulness might play a key role in regulating the state and that some structures might be more recruited by solriamfetol than modafinil or non-pharmacological wakefulness. Our results are particularly relevant for pathologies such as hypersomnia. They open a new era in the study of the mechanisms responsible for inducing wakefulness.

## Introduction

Wakefulness is defined by electroencephalogram (EEG) activation, muscle activity and eye movements. It is well accepted that multiple redundant subcortical structures are responsible for inducing wakefulness (Sulaman et al., 2023). These systems are corresponding to the aminergic (histaminergic, catecholaminergic, serotonergic), cholinergic and the hypocretin/orexin hypothalamic neurons (Sulaman et al., 2023). Although additional populations of wake-inducing neurons have been described recently (Sulaman et al., 2023), their role has not been confirmed. The aim of the present study was to examine over the whole brain all systems potentially inducing wakefulness. Further, we aimed to determine whether wakefulness induced by different drugs such as modafinil (Mod) or solriamfetol (Sol) or by sensory stimulation (non-pharmacological wakefulness, NW) is induced by the same systems. Solriamfetol and modafinil are norepinephrine–dopamine reuptake inhibitors (NDRI) used for the treatment of excessive sleepiness associated with narcolepsy and sleep apnea (Abad & Guilleminault, 2018; Emsellem et al., 2020; Malhotra et al., 2020). Both drugs have been shown to induce wakefulness in mice without the side effects obtained with amphetamine (Hasan et al., 2009). Based on the mode of action of solriamfetol and modafinil, it can be hypothesized that wakefulness induction is due to increased concentrations of norepinephrine and dopamine in the synapses (Baladi et al., 2018). Such increment would lead to enhanced activation of wake-inducing neurons expressing the receptors of one or the two monoamines. Most neurons in the brain express dopamine and/or norepinephrine receptors. Therefore, determining which population(s) of neurons are involved in the induction of waking by solriamfetol and modafinil is challenging. In addition, it has recently been shown that Sol but not Mod has additional agonist activity at the trace amine associated receptor 1 (TAAR1) (Gursahani et al., 2022) and TAAR1 agonists induces W and inhibits NREM sleep (Schwartz et al., 2017). It is therefore possible that the two drugs activate only partly the same wake-inducing structures.

One strategy to identify the neurons responsible for inducing wakefulness would be to determine the effect of solriamfetol, modafinil and NW on neurons already known to be involved in the induction of the state. Another strategy is to identify neurons activated by solriamfetol, modafinil and NW across the entire brain without a priori using the cFos method. Our laboratory has been successfully using the cFos method for more than two decades to identify the neuronal network responsible for inducing paradoxical (REM, Rapid Eye Movement) sleep (PS) and to compare it with wakefulness. Using such a method, we have been the first to identify the neuronal network responsible for inducing muscle atonia during PS and to show that melanin-concentrating hormone neurons (MCH) play a key role in PS generation (Valencia Garcia et al., 2018; Valencia Garcia et al., 2017; Verret et al., 2003). Further, we showed that cortical activation during PS is restricted to a few limbic cortical structures compared to wakefulness opening the avenue to the identification of the function of the state (Renouard et al., 2015).

Hasan et al. (Hasan et al., 2009) qualitatively compared cFos staining obtained after wakefulness induced by solriamfetol, modafinil and amphetamine. Their limited analysis of cFos distribution indicated that the three drugs activate different populations of neurons (Hasan et al., 2009). However, it was not detailed enough to identify the populations of neurons likely responsible for the induction of wakefulness. Further, the cFos method does not allow to determine whether the same or different neurons are activated by different drugs or by wakefulness induced by non-pharmaceutical means.

To reach this objective, we used in the present study an updated genetic method of cFos allowing to determine in the same mice whether the same neurons are activated during two successive periods of wakefulness. We recently used this method based on a new type of transgenic (TRAP) mice (Lee et al., 2020; Maciel et al., 2021; Yamazaki et al., 2021). Using such an innovative model, we did determine whether the same or different neurons are activated by solriamfetol and modafinil or by wakefulness induced by non-pharmaceutical intervention. To this aim, we induced in the same mice one week apart two periods of wakefulness, one using solriamfetol and the other one with modafinil or by putting the mice in an open field. The expression of the reporter gene tdTomato was induced in the activated neurons by injecting 4-hydroxytamoxifen (4-OHT) during the first period of wakefulness. One week later, a second period of wakefulness was induced by another means and the mice were perfused. A whole brain mapping was then made to localize neurons expressing cFos and/or tdTomato. To our knowledge, our report is the first to compare the cFos and the tdTomato staining in the TRAP2 mice over the entire brain. Further, such mapping allowed us to identify neurons activated during wakefulness induced by solriamfetol versus modafinil or non-pharmaceutical induced wakefulness.

## Results

### Sleep-wake analysis

The sleep-wake cycle was analyzed for two hours following the first and second injections of solriamfetol, modafinil or the non-pharmacological wakefulness (NW) induced by sensory stimulation. In the first hour after injections, all animals remained continuously awake, as confirmed by behavioral and electrophysiological recordings (**Fig 1**). In the second hour, wake time started to decrease in particular for modafinil-treated mice for the first session (H[2,13] = 6.40, *p* = 0.041, third 30-minute interval; H[2,13] = 8.31, *p* = 0.016, fourth 30-minutes interval; Kruskal-Wallis test) (**Fig 1b**). In the second experimental session, a significant decrease of wake time was observed both for Sol- and Mod-treated groups but only during the fourth 30-minute interval (H[2,13] = 7.48, *p* = 0.024, Kruskal-Wallis test, **Fig 1c**).

**Fig 1.**
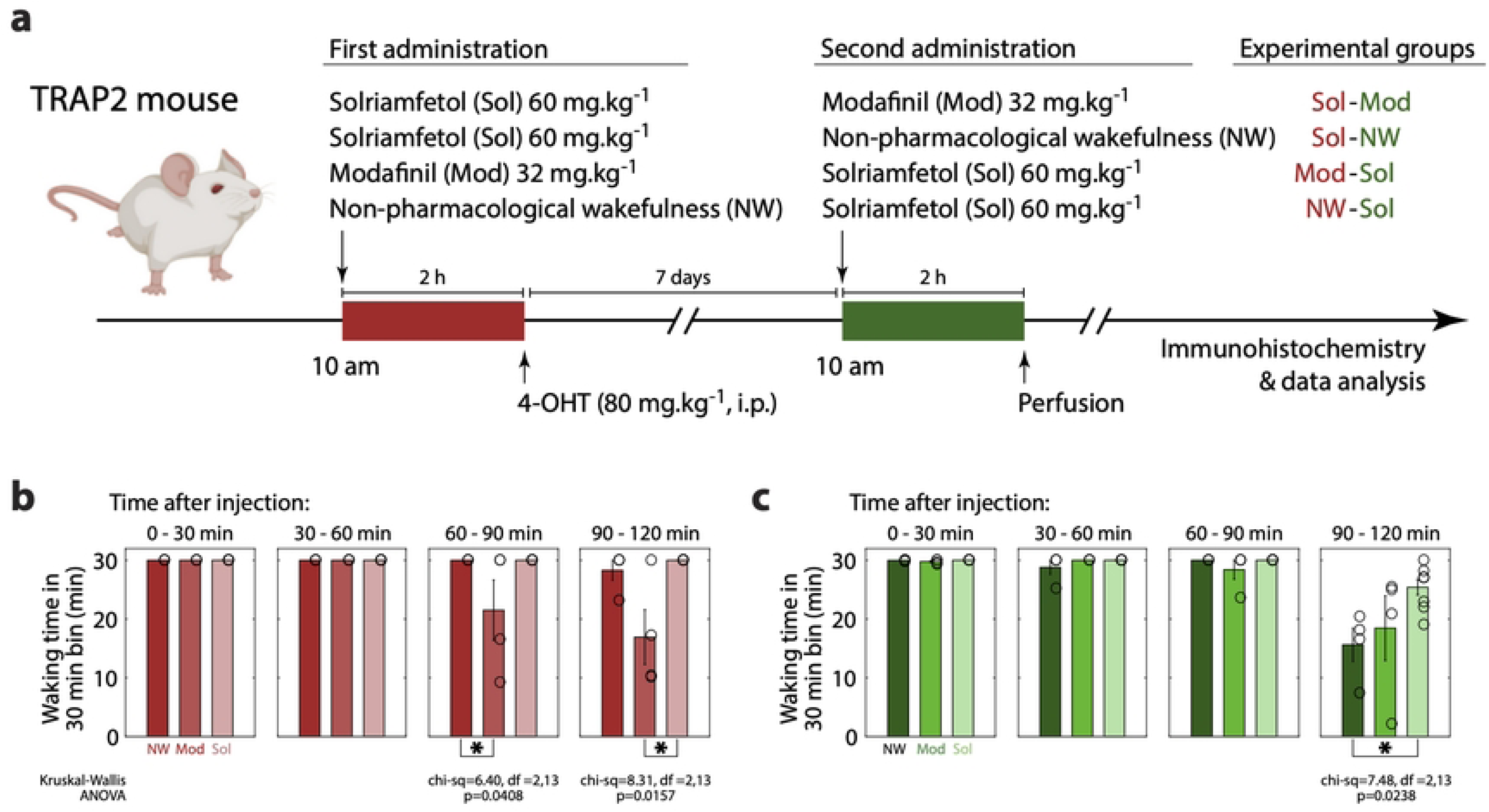
Experimental design and sleep recordings. **(a)** Male adult (8-12 weeks old mice) TRAP2 mice were placed in the recording chamber for seven days before the beginning of the experiments. On the first experimental day, animals received an IP injection of solriamfetol (Sol) or modafinil (Mod) at 10 a.m. The introduction of new objects in the recording chamber and occasional sensory stimulation was used to induce non-pharmacological wake (NW). All animals were injected with 80 mg/kg of 4-OHT 2 hours after the administration of the experimental drugs or wake induction to induce the expression of the reporter gene (tdTomato) in cFos-expressing neurons. Seven days after the first condition, animals returned to the recording chamber and received an administration of the other drug or were subjected to NW. Animals were perfused 2 hours later and their brains were processed for immunofluorescence staining of cFos. Each group (top-right) contained four animals (16 animals in total). **(b-c)** Time spent in wakefulness in the first (b) and second (c) experimental session per 30 minutes, starting at the injection of solriamfetol (Sol), modafinil (Mod) or the beginning of the induction of NW, open field). Nonparametric Kruskal-Wallis followed by post hoc Mann-Whitney test, * p<0.05.

### Overall analysis

To determine the structures differentially activated by Sol, Mod and NW as well as those strongly activated during all conditions, the number of tdT, cFos and double-labeled neurons was quantified in the whole brain in all mice (10 sections for each 16 mice). We evaluated 164 structures of interest (SOI) without priori assumptions, of which 109 structures met the inclusion criteria (values from at least three animals per group), organized in eight macroregions: cortex, telencephalon, thalamus, hypothalamus, hippocampal formation, mesencephalon, pons and medulla. A total of 205,316 tdT+ cells and 217,039 cFos+ cells were counted in all mice and no statistical difference was observed between the number of tdT and cFos positive cells (mean±SEM per mouse, tdT: 12,832±1,462 and cFos: 13,565±1,354, p=0.57, paired t-test). Moreover, the expression of tdT and cFos was strongly correlated in the structures analyzed in all animals (R^2^=0.76±0.02, mean±SEM, N=16; p<10^-20^, Pearson correlation on log10 transformed number of tdT and cFos positive cells; **Fig 2**).

**Fig 2.**
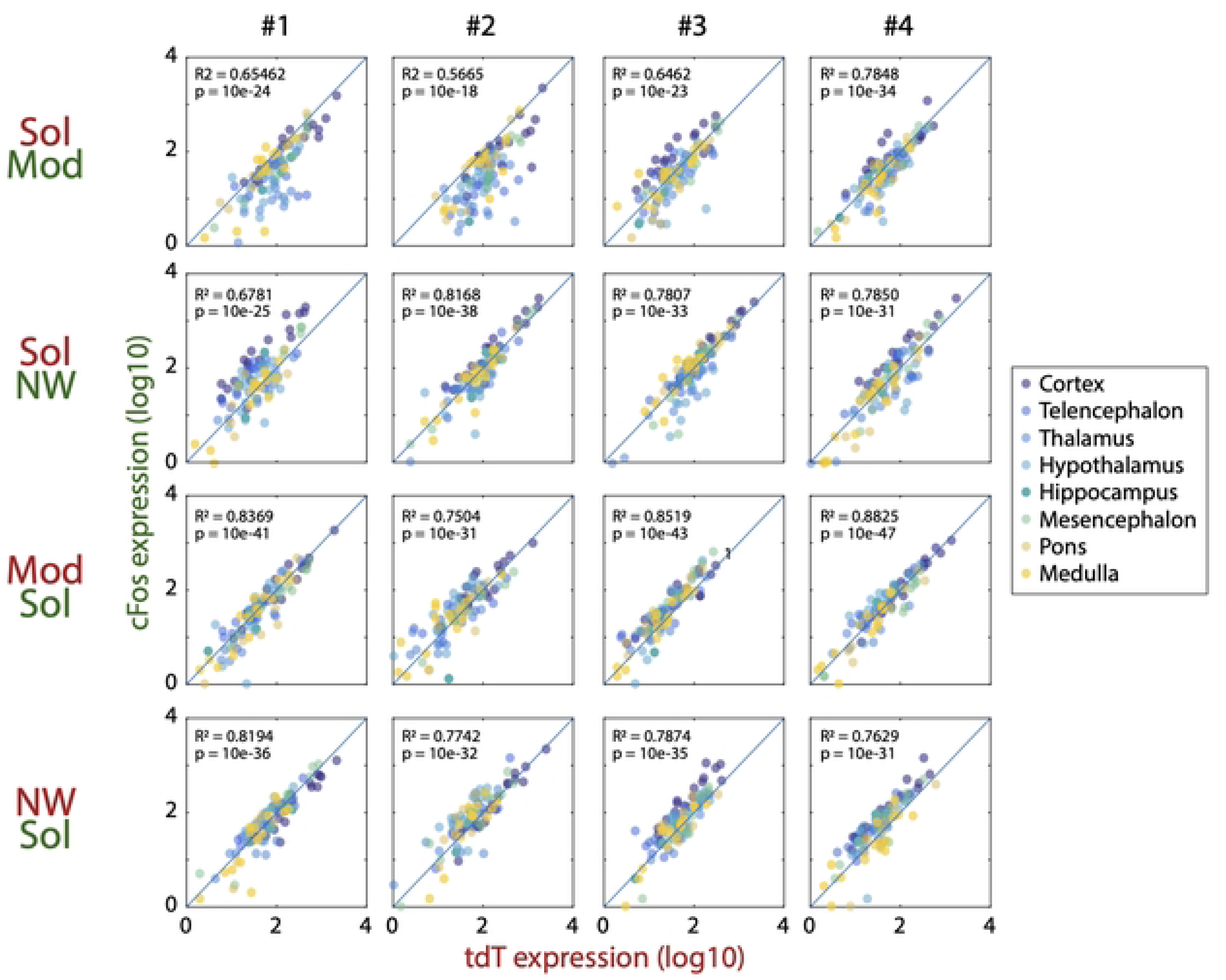
Correlation between the expression of tdTomato and cFos. The log transforms numbers of tdT and cFos labeled neurons per structure are linearly correlated in the four animals per group (left to right, Pearson correlation, R2 and p-value shown in each panel). Note that most of the structures are closed to the diagonal line.

We then calculated the density of tdT and cFos-labeled neurons across brain regions and individual structures and found an overall similar pattern in cell distribution for the two markers (**Fig 3**). The highest cell density was found in the hypothalamus, in the paraventricular hypothalamic nucleus (PVN) and the supraoptic nucleus (SON), while the hippocampal formation (HF) showed the lowest cell density across all brain regions (**Fig 3**).

**Fig 3.**
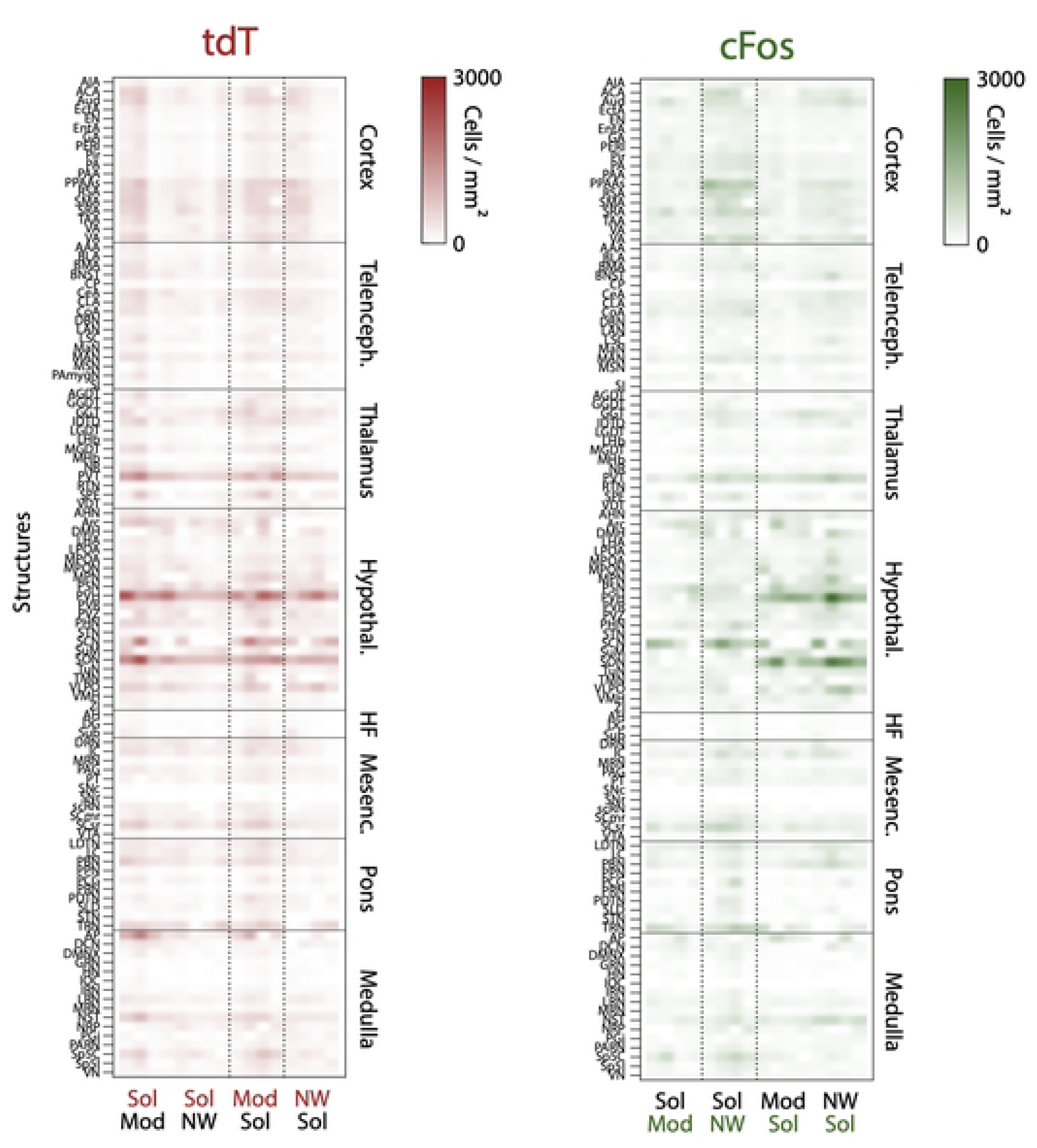
Brain-wide expression of tdTomato (tdT) and cFos after solriamfetol (Sol), modafinil (Mod), and non-pharmacological wakefulness (NW). Cell density heatmap representation of tdTomato (left) and cFos (right) in the cortex, telencephalon, thalamus, hypothalamus, hippocampal formation (HF), mesencephalon, pons, and medulla. Each column represents one mouse. The cell density is color coded from zero to 3000 cells/mm2.

To identify structures differentially affected by Sol, Mod, or NW, a two-way ANOVA was performed on tdT+ and cFos+ cell densities across all structures. The statistical analysis showed that 20 structures were more activated in the Sol condition **(Table 1, Supp. Table 1)** than in the two other conditions. Five structures were more activated both by NW and SOL compared to Mod **(Supp. Table 2)**. Finally, 19 structures were more activated by NW than the two other conditions **(Supp. Table 3)**. Surprisingly, no structure was more activated by Mod than the two other conditions. Finally, 65 structures were not differentially activated in the three conditions.

**Table 1.** Structures modified by solriamfetol. Columns show from the left to the right, the two-way ANOVA p-values (factor 1: experimental treatment; factor 2: marker, and interaction between the two factors), the mean values of tdT and cFos cell densities per mm2 and the standard error of the mean (SEM) for each condition and structure. The stars show p<0.05 (Tukey-Kramer post-hoc pairwise comparison) within each marker. (*) *vs* NW, and (**) *vs* Sol groups.

| Structure | ANOVA p-value |  |  | tdT |  |  | cFos |  |  |
| --- | --- | --- | --- | --- | --- | --- | --- | --- | --- |
|  | Fact.1 | Fact.2 | Inter. | Sol | Mod | NW | Sol | Mod | NW |
| Area postrema | 0.0177 | 0.2265 | 0.5224 | 851.9 ± 169.4 | 252.5 ± 74.4 | 304.0 ± 52.0 | 454.0 ± 90.8 | 179.0 ± 109.2 | 183.0 ± 3.5 |
| Bed nuclei of the stria terminalis | 0.0041 | 0.4225 | 0.6200 | 273.0 ± 34.8 | 144.5 ± 14.2 | 172.3 ± 22.5 | 288.4 ± 71.3 | 59.5 ± 29.4 ** | 131.9 ± 24.4 |
| Central amygdalar nucleus | 0.0001 | 0.3596 | 0.8382 | 346.6 ± 40.4 | 154.5 ± 19.2** | 198.8 ± 24.6 | 294.0 ± 39.1 | 126.3 ± 10.2 ** | 189.5 ± 18.8 |
| Lateral reticular nucleus | 0.0565 | 0.6556 | 0.1631 | 200.6 ± 21.0 | 111.2 ± 5.7 ** | 184.5 ± 25.8 | 168.2 ± 9.6 | 154.1 ± 14.5 | 150.4 ± 37.7 |
| Lateral septum | **0.0363** | 0.4397 | 0.8726 | 217.0 ± 48.1 | 98.6 ± 23.1 | 129.8 ± 33.7 | 177.0 ± 48.2 | 41.6 ± 24.8 | 129.5 ± 20.8 |
| Medial preoptic area | **0.0355** | 0.4481 | 0.8580 | 246.3 ± 23.8 | 127.1 ± 30.5 | 212.8 ± 45.0 | 308.8 ± 76.4 | 129.5 ± 18.7 | 255.5 ± 25.9 |
| Median preoptic nucleus | **0.0091** | 0.7702 | 0.4584 | 396.1 ± 70.4 | 141.7 ± 23.1 | 338.0 ± 61.8 | 513.7 ± 126.7 | 40.1 ± 8.9 | 242.5 ± 29.2 |
| Nucleus of the solitary tract | **0.0036** | 0.7859 | 0.8597 | 495.0 ± 75.6 | 244.0 ± 19.7 | 290.8 ± 13.8 | 490.5 ± 66.7 | 232.0 ± 53.6 | 358.3 ± 94.6 |
| Parabrachial nucleus | **0.0150** | 0.9650 | 0.8463 | 409.4 ± 55.8 | 282.5 ± 67.4 | 209.3 ± 23.2 | 409.4 ± 63.6 | 243.8 ± 47.3 | 255.3 ± 75.7 |
| Paraventricular hypothalamic n. | **0.0000** | **0.0161** | 0.1067 | 1603.8 ± 159.8 | 874.3 ± 28.5 | 1178.3 ± 227.1 | 1632.5 ± 265.1* | 271.1 ± 125.1** | 345.8 ± 50.6 |
| Paraventricular n. of the thalamus | **0.0008** | 0.1227 | 0.2397 | 988.5 ± 116.1 | 504.0 ± 27.5* | 609.8 ± 49.3 | 696.3 ± 65.4 | 395.8 ± 94.6 | 628.3 ± 66.0 |
| Periventricular region | **0.0070** | 0.6684 | 0.6872 | 434.3 ± 59.1 | 206.5 ± 28.8 | 183.5 ± 45.4 | 360.0 ± 91.7 | 150.8 ± 38.1 | 232.8 ± 15.2 |
| Supraoptic nucleus | **0.0000** | **0.0185** | **0.0130** | 1358.9 ± 177.9 | 824.5 ± 24.9 | 958.3 ± 73.3 | 1619.6 ± 249.9* | 111.6 ± 51.1** | 132.0 ± 27.3 |
| Tuberal nucleus | **0.0058** | 0.1978 | 0.2706 | 224.1 ± 39.1 | 151.5 ± 26.7 | 189.0 ± 23.7 | 334.8 ± 36.3 | 151.8 ± 28.2** | 207.0 ± 33.5 |

The values of the reactivation index (RI) from 109 structures and 16 animals are shown in **Fig 4**. The histogram is well described by a gamma distribution (R2=0.92, p=1.5×10^-18^) with average RI of 0.26 and median RI of 0.24 (considering all measurements). Irrespective of the experimental condition, the average RI was higher in the hypothalamus (RI=0.314±0.010), mesencephalon (RI=0.318±0.012), pons (RI=0.315±0.014), and medulla (RI=0.289±0.010) compared with cortex (RI=0.198±0.005), telencephalon (RI=0.225±0.008) and thalamus (RI=0.209±0.011; F_[7,1657]_=34.07, p=1.4×10^-44^, one-way ANOVA). The hippocampal formation showed the lowest RI among macrostructures (RI=0.079±0.007).

**Fig 4.**
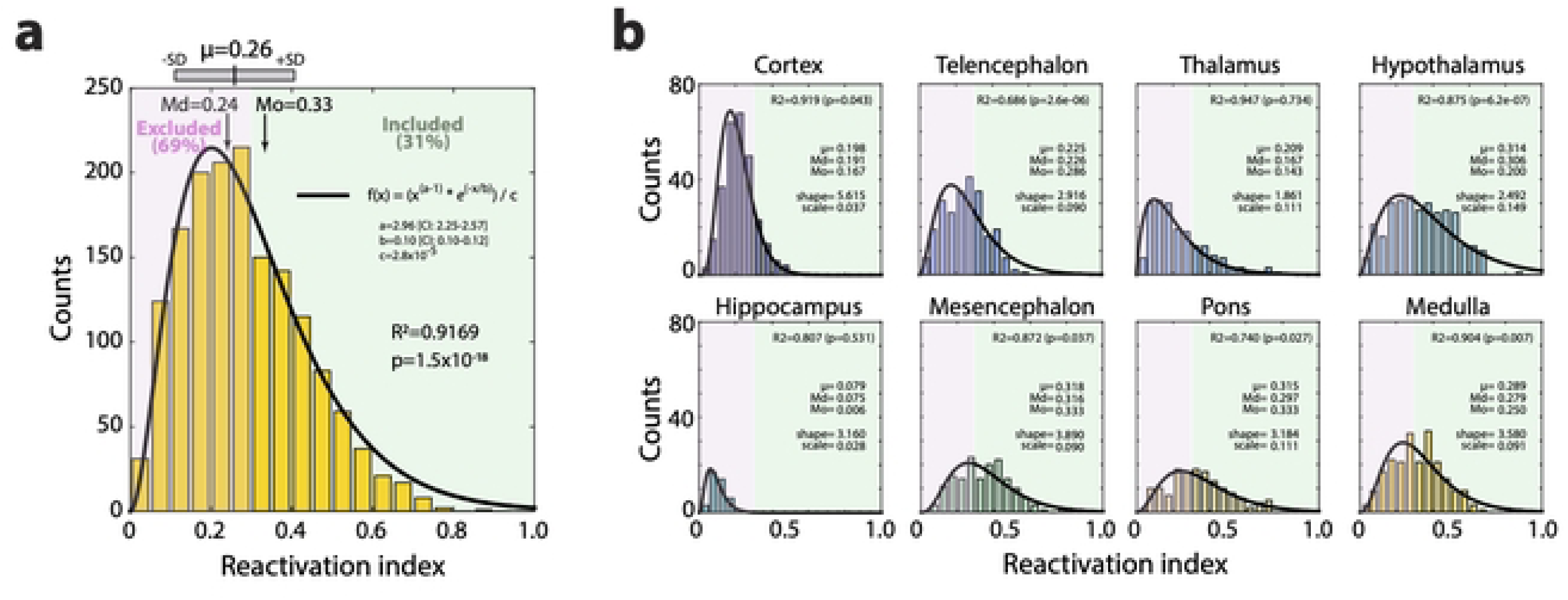
Distribution of the reactivation index (RI). (**a**) Histogram of the RI for 109 structures and 16 animals (bin size=0.05). The histogram was fitted to a gamma distribution with parameters shape (a=2.96) and scale (b=0.10; R2=0.91, p=1.5×10^-18^). The mean (µ), median (Md) and mode (Mo) is shown in the Fig, together with the parameters (CI: confidence interval), R2 and p-value. (**b**) Histograms of the RI for each macroregion. A threshold above 0.3 in the four groups was set as an inclusion criteria for populations of neurons activated during all types of wakefulness.

To evaluate whether the RI was affected by the experimental conditions, we performed a one-way ANOVA for each structure. Twenty-eight out of 109 structures (26%) showed p-values < 0.05 indicating differential activation (**Fig 5**). Most statistically significant differences were found in the cortex (12/18), telencephalon (5/16) and thalamus (4/13), followed by the hypothalamus (4/22), medulla oblongata (2/16) and pons (1/10). The mesencephalon (11 structures) and the hippocampal formation (3 structures) showed no significant difference in RI between conditions (**Fig 5a**). Post hoc (unpaired t test) analysis revealed that 14 structures (11 cortical) had significantly higher RI in the Sol-NW mice compared to the three other groups of mice (**Fig 5b**). In contrast, 10 subcortical structures exhibited higher reactivation in the Mod-Sol and/or the NW-Sol conditions compared to the Sol-Mod and/or the Sol-NW conditions (**Fig 5b**). Additionally, three thalamic structures displayed higher RI in the NW-Sol group than in the Sol-Mod group (**Fig 5b**). Finally, a group of 11 structures displayed an RI above 0.3 across all groups of mice (**Table 3**). In the next section, we detail the significant differences revealed by the cell densities and reactivation analyses for each structure.

**Fig 5.**
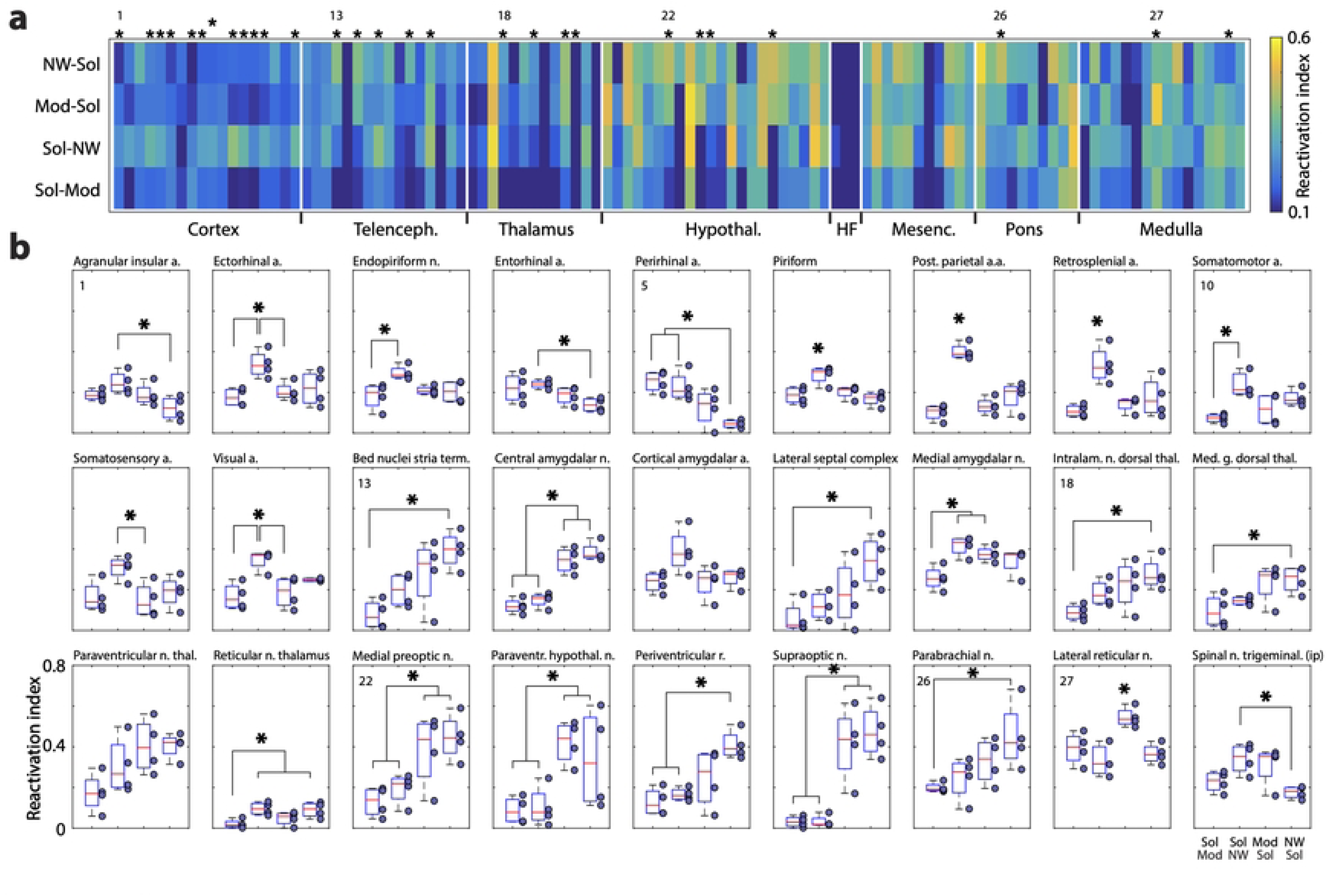
Reactivation index. (**a**) Heatmap of the averaged reactivation index (RI) for the four experimental groups (Y-axis). Asterisks on top of the panel represent structures with p<0.05 (one-way ANOVA). The first statistically different structure in each macroregion is labeled with its number starting by one. (**b**) Boxplots of the structures with p<0.05 in one-way ANOVA (shown with asterisks in a and b) starting by the cortex and then the subcortical structures. The numbers correspond to those shown in a. A RI=1 means that all tdT cells also express cFos, while RI=0 means that no tdT cell co-expresses cFos. Note that for the cortical amygdalar nucleus and the paraventricular thalamic nucleus, the difference between conditions did not reach statistical difference when using the Tukey-Kramer post-hoc test.

### Structures differentially activated by Sol, Mod or NW

To uncover the brain structures with significantly different patterns of activation across the experimental conditions, we first performed a two-way ANOVA on the density of tdT+ and cFos+ neurons in the Sol, Mod and NW conditions. Such statistical analysis did not take advantage of the model of the TRAP mice. To do so, we also realized paired t tests to compare the Tdt and cFos density during two conditions in the same mice. Then, a one-way ANOVA was performed on the reactivation index (RI), calculated as the ratio of tdT-cFos double-labeled cells to the total number of tdT+ cells. Post-hoc analyses were performed to identify specific group differences. Such a multiple statistical strategy allowed us to be sure that some structures were differentially activated by one of the two conditions. Below, we provide a detailed description of our analysis.

### Identification of the structures specifically activated by Sol

The two-way ANOVA analysis first showed that a group of 14 structures was characterized by a significantly increased density of tdT and cFos activated cells in the Sol condition compared to the Mod and NW conditions (**Table 1**). This group was composed of lateral septum, bed nucleus of the stria terminalis **(Fig 6)**, medial and median preoptic nucleus, central amygdalar nucleus **(Fig 6)**, paraventricular thalamic **(Fig 9)** and hypothalamic nuclei **(Fig 6)**, supraoptic nucleus **(Fig 6)**, periventricular region, tuberal nucleus, lateral parabrachial nucleus **(Fig 7)**, lateral reticular nucleus **(Fig 7)**, nucleus of the solitary tract **(Fig 7)** and area postrema. The Tukey-Kramer post-hoc confirmed only partly the differences displayed by the two-way ANOVA which was expected since it corrects for multiple comparisons and is therefore quite conservative. It showed that the density of tdT and cFos neurons was significantly higher in the Sol condition compared to the Mod one in the central amygdalar nucleus. For the bed nucleus of the stria terminalis and the tuberal nucleus, only the density of cFos neurons was higher in the Sol condition compared to the Mod one. Importantly, the density of cFos+ neurons but not that of tdT was significantly higher in the Sol condition compared to the Mod and NW conditions in the supraoptic and the paraventricular hypothalamic nuclei. These were the only structures for which the density of tdT labeled neurons was significantly higher than that of cFos for the Mod and NW conditions. Such a difference between markers is clearly visible in **Fig 6c,e**. Finally, the density of tdT neurons was significantly lower in the Mod condition compared to the NW or the Sol conditions in the paraventricular nucleus of the thalamus and lateral reticular nucleus, respectively.

**Fig 6.**
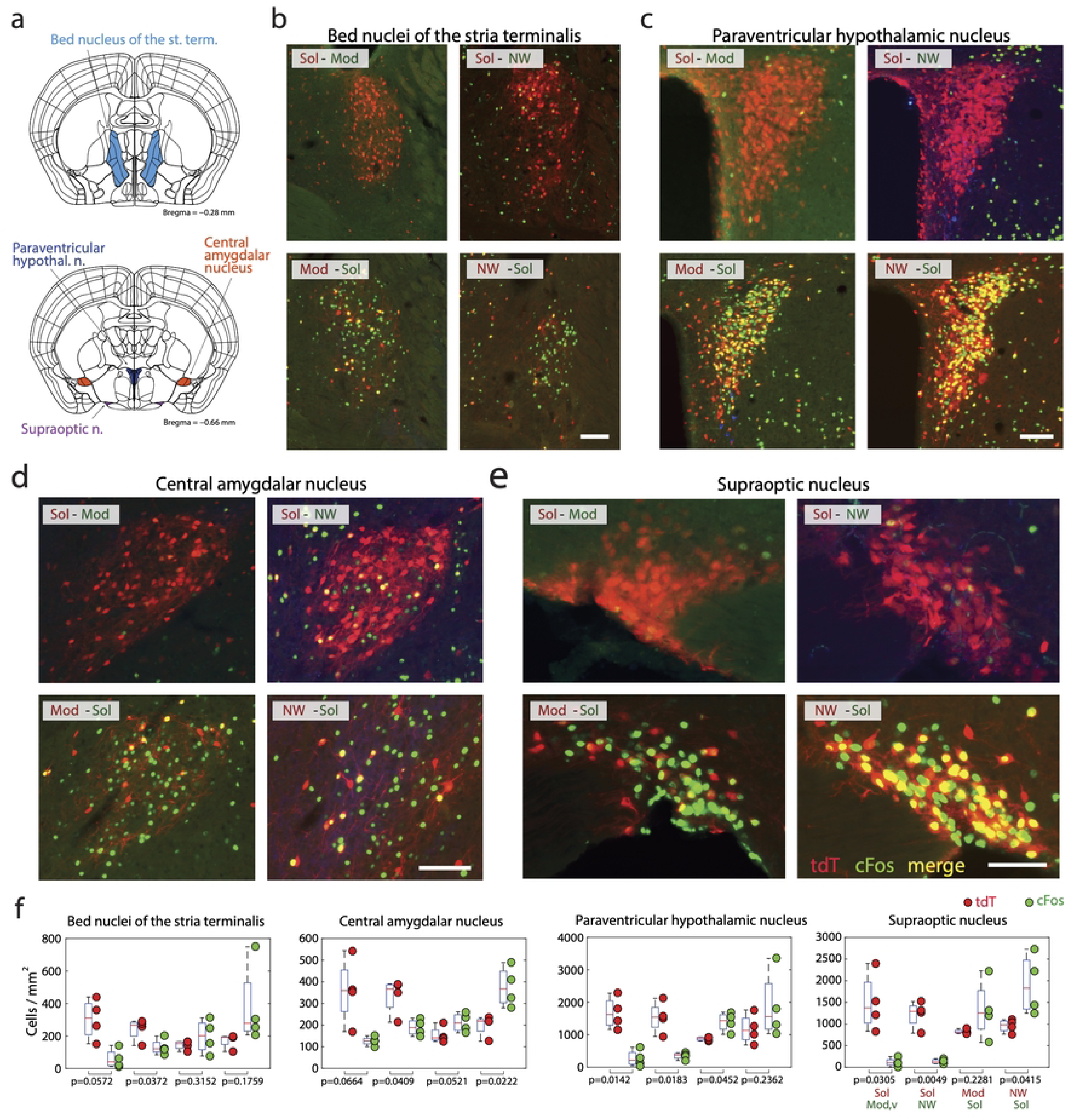
Forebrain structures are more activated by Solriamfetol. (**a**) Drawings showing the localization of the structures (**b-e**) Photomicrographs illustrating the distribution of tdT (red), cFos (green) and double-labeled neurons (yellow) in one representative mouse per experimental group in four diencephalic structures. Note that for the four structures, more cFos or tdT neurons are labeled in the Sol condition compared to the Mod and NW conditions (**f**) Boxplots showing the density (cell/mm^2^) of tdT+ (red) and cFos+(green) neurons in each mice of the four groups (Sol-Mod, Sol-NW, Mod-Sol, NW-Sol, n = 4 per group). Paired t tests between tdT and cFos densities in the same mice, p < 0.05*. Boxes, median; dots, individual mouse values; boxes, first and last quartiles; whiskers, minimum and maximum values excluding outliers. Scale b,c,d: 100 μm; scale e: 60 μm.

**Fig 7.**
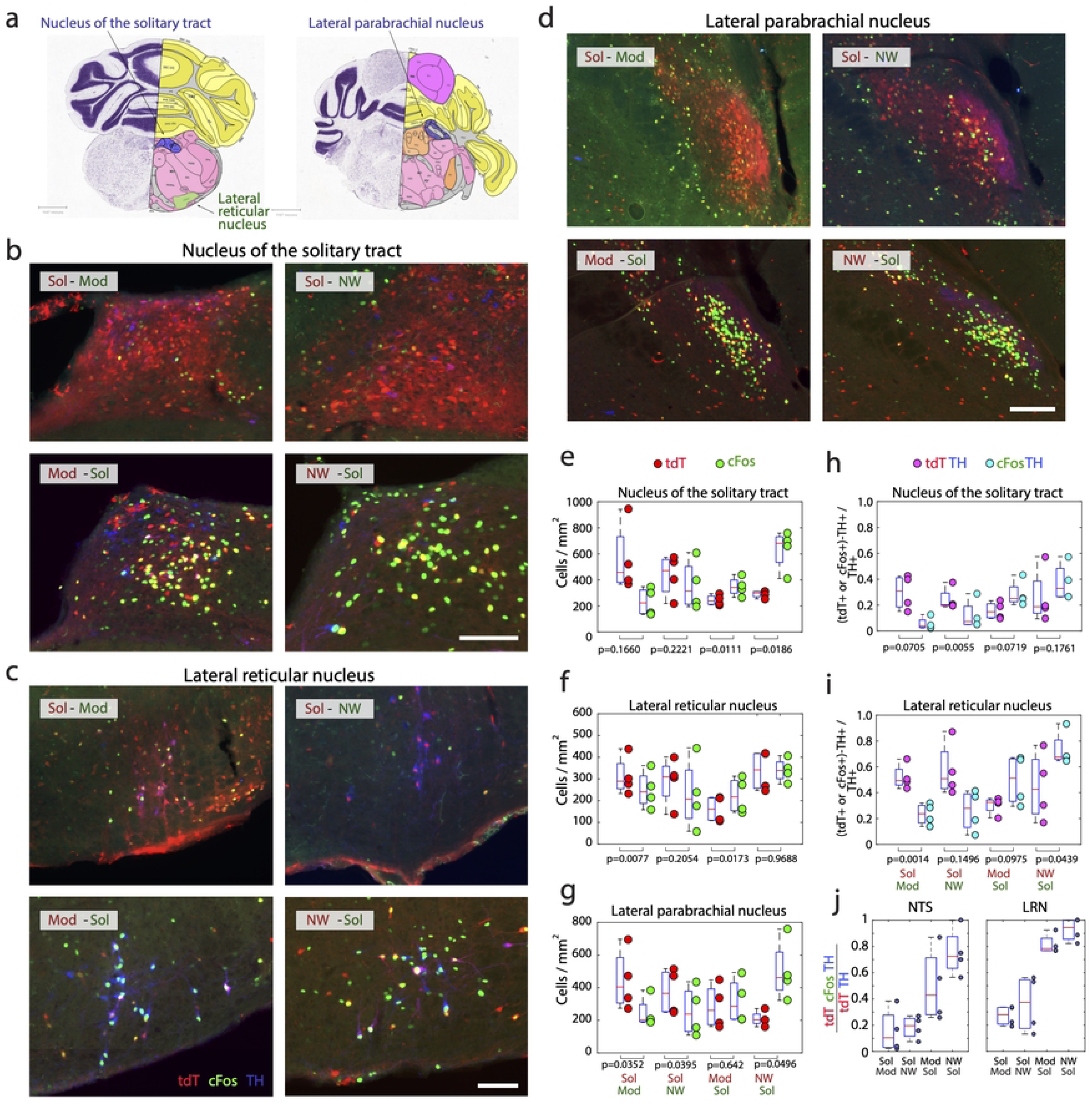
Ponto-medullary structures are more activated by Solriamfetol. (**a**) Drawings showing the localization of the structures. (**b-d**) Photomicrographs illustrating the distribution of tdT (red), cFos (green), TH (blue), tdT and TH double-labeled neurons (purple) and, cFos and TH double labeled neurons (white nuclei) in one representative mouse per experimental group in one pontine and two medullary structures. Note that for the three structures, more cFos or tdT neurons are labeled in the Sol condition compared to the Mod and NW conditions. (**e-g**) Boxplots showing the density (cell/mm^3^) of tdT+ (red) and cFos+ (green) neurons in each of the four groups (Sol-Mod, Sol-NW, Mod-Sol, NW-Sol, n = 4 per group). Paired t test when comparing tdT and cFos for each group. **(h-i)** Ratio of TH neurons double-labeled with tdT or cFos in the nucleus of the solitary tract (NTS) and lateral reticular nucleus (LRN). Note the significantly higher level of activation of the TH neurons in the Sol condition compared to the two other conditions. **(j)** Reactivation index (tdTomato-cFos-TH+ neurons over total tdTomato-TH+ neurons) for LRN and NTS. Significance one-way Anova and Post-hoc test. p < 0.05*. Boxes, median; dots, individual mouse values; boxes, first and last quartiles; whiskers, minimum and maximum values excluding outliers. Scale b,c: 100 μm; scale d: 200 μm.

The comparison between tdT and cFos densities in the two conditions applied to the same mice using paired t tests was globally in line with the above analysis. The density of Tdt neurons in the central amygdalar nucleus was indeed significantly superior to that of cFos in the Sol-Mod and Sol-NW groups while that of cFos was superior to tdT in the NW-Sol group **(Fig 6f)**. It was less clear in the bed nucleus of the stria terminalis in which tdT density was significantly higher than for cFos only in the Sol-NW group **(Fig 6f)**. In the tuberal nucleus, cFos density was significantly higher to tdT in the NW-Sol group while the opposite occurred in the Sol-Mod group (not illustrated). The supraoptic, paraventricular and lateral parabrachial nuclei behave quite similarly with significantly higher density of tdT neurons compared to cFos in the mice with Sol as the first condition and the opposite with mice with Sol as the second conditions excepting for the Mod-Sol group for the supraoptic **(Fig 6f)** and lateral parabrachial **(Fig 7g)** nuclei and the NW-Sol mice for the paraventricular hypothalamic nucleus **(Fig 6f).** For the medial preoptic area, significantly higher tdT neurons density than cFos was found in the Sol-Mod group and conversely more cFos than tdT in the Mod-Sol group while no difference was observed in the Sol-NW and NW-Sol group indicating a higher activation during Sol than Mod but not compared to NW (not illustrated). For the nucleus of the solitary tract and the lateral reticular nucleus **(Fig 7e,f)**, a significantly higher density of cFos compared to tdT was observed in the Mod-Sol group and in the NW-Sol group specifically for the nucleus of the solitary tract. In the periventricular region, there was significantly higher tdT density than cFos only in the Sol-Mod group. For the lateral septal complex, median preoptic nucleus and the area postrema, no difference in tdT and cFos density was observed in the four groups of mice (not shown).

When examining the reactivation index, we found that all of the structures identified above excepting the median preoptic nucleus, the tuberal nucleus, the nucleus of the solitary tract and the area postrema displayed a significantly different reactivation index between at least two groups **(Figs 5-7)**. The first category of structures composed of the central amygdalar nucleus, the medial preoptic area, the paraventricular hypothalamic and supraoptic nuclei showed a significantly higher RI in the Mod-Sol and NW-Sol conditions compared to the Sol-Mod and Sol-NW conditions. The second category of structures composed of the bed nucleus of the stria terminalis, lateral septal complex, periventricular region and lateral parabrachial nucleus showed a significantly higher RI in the NW-Sol condition compared to the Sol-Mod condition. Finally, the lateral reticular nucleus showed a higher RI in the Mod-Sol condition compared to the three other conditions.

### Structures containing a higher density of tdT but not cFos neurons in Sol condition

The second group of structures was composed of thalamic nuclei showing an increased number of tdT cells in the Sol condition compared to the NW and Mod conditions as shown by two-way ANOVA analysis (**Supp. Table 1).** Surprisingly, these structures did not show an increased number of cFos-positive neurons in the Sol condition compared to the NW and Mod conditions. Significativity was confirmed by the Tukey-Cramer test for the anterior group of the dorsal thalamus, the lateral and medial group of the dorsal thalamus and the reticular nucleus of the thalamus but not for the nucleus reuniens and the intralaminar nuclei of the dorsal thalamus **(Supp. Table 1)**. Then, a paired t test was performed to determine whether these structures show a significantly higher number of tdT than cFos neurons in the Sol-NW and/or the Sol-Mod conditions and not conversely in the NW-Sol and/or Mod-Sol conditions. The anterior group of the dorsal thalamus, lateral and ventral group of the dorsal thalamus, reticular nucleus of the thalamus showed indeed a higher density of tdT than cFos in the Sol-NW and/or the Sol-Mod conditions. Three of these thalamic structures, namely the intralaminar nuclei of the dorsal thalamus, medial group of the dorsal thalamus and the reticular nucleus of the thalamus showed a significantly higher RI in the NW-Sol condition compared to the Sol-Mod condition (**Fig 5**). Altogether, these results clearly indicate that some thalamic nuclei show a higher density of tdT but not cFos neurons in the Sol than in the Mod and NW conditions.

**Supp. Table 1.** Structures with higher tdT density in Sol condition. Columns show from the left to the right, the two way ANOVA p-values (Factor 1, treatment; Factor 2, staining method; and interaction between the two), the mean values of tdT and cFos cell densities per mm2 and the standard error of the mean (SEM) or each condition and structure. The stars show p<0.05 (Tukey-Kramer post-hoc pairwise comparison) within marker. (**) *vs* Sol-tdT.

| Structure | ANOVA p-value |  |  | tdT |  |  | cFos |  |  |  |  |  |
| --- | --- | --- | --- | --- | --- | --- | --- | --- | --- | --- | --- | --- |
|  | Fact.1 | Fact.2 | Inter. | Sol | Mod | NW | Sol | Mod | NW | Sol | Mod | NW |
| Anterior group of the | 0.0296 | 0.1449 | 0.0980 | 245.1 ± 52.6 | 52.3 ± 32.8** | 114.8 ± 32.9 | 85.4 ± 28.6** | 34.6 ± 25.6** | 126.0 ± 40.5 |  |  |  |
| Intralaminar nuclei of | 0.0253 | 0.1243 | 0.1382 | 377.1 ± 56.3 | 178.4 ± 44.3 | 205.7 ± 41.5 | 203.2 ± 42.9 | 102.5 ± 30.6** | 243.5 ± 53.4 |  |  |  |
| Lateral group of the | 0.0495 | 0.0353 | 0.0031 | 127.3 ± 22.3 | 42.8 ± 9.4** | 51.8 ± 4.7** | 32.1 ± 4.1** | 37.0 ± 16.2** | 62.5 ± 9.6 |  |  |  |
| Medial group of the | 0.0104 | 0.0620 | 0.1165 | 303.4 ± 52.9 | 127.8 ± 15.1** | 129.6 ± 39.6** | 145.1 ± 22.9** | 77.9 ± 35.1** | 135.5 ± 10.7 |  |  |  |
| Nucleus of reuniens | 0.0231 | 0.0720 | 0.0369 | 310.8 ± 61.5 | 131.5 ± 18.5 | 123.7 ± 24.5 | 125.7 ± 19.4** | 53.3 ± 20.5** | 175.9 ± 34.5 |  |  |  |
| Reticular nucleus of | 0.0211 | 0.1439 | 0.0159 | 64.9 ± 14.2 | 20.2 ± 5.6** | 26.0 ± 2.1 | 21.4 ± 4.1** | 6.0 ± 2.5** | 44.6 ± 10.1 |  |  |  |

### Structures containing a higher density of tdT and cFos neurons in Sol and NW conditions

In a third group of 5 structures namely the anterior hypothalamic nucleus, diagonal band nucleus, lateral habenula, lateral hypothalamic area **(Fig 10d,f)** and zona incerta, the two way ANOVA showed that there was statistical differences between conditions and for three of them also between markers **(Sup. Table 2)**. Although there were more tdT and cFos neurons in the NW and Sol conditions compared to the Mod condition in the 5 structures, the Tukey Kramer test showed that cFos density was significantly higher only during NW compared to Mod in the anterior hypothalamic nucleus and the lateral hypothalamic area **(Sup. Table 2)**. Then, statistical differences were determined using a paired t test between two conditions in the same mice. In the anterior hypothalamic nucleus, there was a significantly higher density of cFos than tdT neurons in the Mod-Sol and NW-Sol groups. In the diagonal band nucleus, the density of cFos was significantly higher than tdT in the Mod-Sol, Sol-NW but also NW-Sol conditions. In the lateral habenula, the density of tdT was higher than cFos only in the Sol-Mod group. In the lateral hypothalamic area, the density of cFos was higher in the Sol-NW and the NW-Sol conditions. For the zona incerta, no difference was observed. Finally, the RI was not different between groups for these structures and was not above 0.3 for all conditions although it was very close to it for the lateral hypothalamic area **(Fig 10f).** Altogether, these results suggest that the modifications between conditions are marginal in these five structures.

**Supp. Table 2.** Structures modified by NW and solriamfetol. Columns show from the left to the right, the two way ANOVA p-values (Factor 1, treatment, Factor 2, marker and interaction between the two), the mean values of tdT and cFos cell densities per mm2 and the Standard error of the mean (SEM) for each condition and structure. The stars show p<0.05 (Tukey-Kramer post-hoc pairwise comparison) within each. (*) *vs* NW, and (**) *vs* Sol within marker; (#) *vs* NW-cFos, and (&) *vs* Sol-cFos between markers.

| Structure | ANOVA p-value |  |  | tdT |  |  | cFos |  |  |  |  |  |
| --- | --- | --- | --- | --- | --- | --- | --- | --- | --- | --- | --- | --- |
|  | Fact.1 | Fact.2 | Inter. | Sol | Mod | NW | Sol | Mod | NW | Sol | Mod | NW |
| Anterior hypothalamic | 0.0077 | 0.0114 | 0.0685 | 257.1 ± 40.4 | 144.5 ± 13.3 | 131.5 ± 20.2 | 324.5 ± 40.0 | 145.4 ± 36.7* | 361.8 ± 61.8 |  |  |  |
| Diagonal band | 0.0046 | 0.0007 | 0.6876 | 93.6 ± 13.2& | 49.2 ± 16.8&# | 110.6 ± 26.2 | 171.2 ± 19.5 | 93.3 ± 23.0 | 183.0 ± 18.3 |  |  |  |
| Lateral habenula | 0.0374 | 0.6891 | 0.4073 | 100.8 ± 16.4 | 64.4 ± 16.5 | 66.2 ± 16.8 | 93.7 ± 17.6 | 29.2 ± 7.6 | 89.2 ± 25.1 |  |  |  |
| Lateral hypothalamic | 0.0086 | 0.0236 | 0.0470 | 174.4 ± 22.0 | 100.9 ± 20.6# | 130.0 ± 23.1# | 177.6 ± 19.3 | 118.7 ± 18.1* | 248.0 ± 22.3 |  |  |  |
| Zona incerta | 0.0382 | 0.3720 | 0.3526 | 147.2 ± 24.0 | 84.7 ± 18.6 | 125.2 ± 18.8 | 150.0 ± 32.2 | 72.3 ± 8.1 | 205.3 ± 46.5 |  |  |  |

### Structures containing a higher density of cFos neurons in NW condition

A group of 19 structures did show a significantly higher density of cFos-labeled cells specifically in the NW condition compared to Sol and/or Mod conditions (**Supp. Table 3**). However, these structures did not show a significantly higher density of tdT than cFos neurons in the NW condition compared to the Sol and Mod ones. Nine of these structures were cortical namely the Ammon’s horn, anterior cingulate area, ectorhinal and entorhinal areas, perirhinal area, piriform area, posterior parietal association area, retrosplenial area and visual area **(Supp. Fig 1)**. Five of these structures were located in the forebrain namely the basomedial and medial amygdalar nuclei, caudoputamen, magnocellular nucleus, substantia innominata and subthalamic nucleus. Finally, four of these structures were located in the brainstem namely the midbrain reticular nucleus, pedunculopontine nucleus, pontine reticular nucleus and superior colliculus, motor related. Then, we examined using a paired t test whether some structures showed a significantly higher density of cFos than tdT neurons in the same mice in the Sol-NW group and not conversely in the NW-Sol group. There were significantly more cFos than tdT neurons in the Sol-NW group but not in the NW-Sol group in 9 of the structures identified above namely the Ammon’s horn, anterior cingulate area, ectorhinal area, piriform area, medial amygdalar nucleus, magnocellular nucleus, midbrain reticular nucleus, pontine reticular nucleus, superior colliculus motor related. Nine additional structures also showed such statistical differences namely agranular insular area, claustrum, cortical amygdalar area, diagonal band nucleus, endopiriform nucleus, inferior colliculus, lateral preoptic area, somatosensory areas and tegmental reticular nucleus. Finally, we determined the structures with a higher reactivation index in the Sol-NW group compared to at least one of the other three groups of mice **(Fig 5)**. Fourteen of the structures identified above showed a higher reactivation index specifically in the Sol-NW group namely the ectorhinal area, piriform area, entorhinal areas, perirhinal area, posterior parietal association area, retrosplenial area, visual area, medial amygdalar nucleus, spinal trigeminal nucleus, interpolar part, agranular insular area, somatomotor and somatosensory areas, cortical amygdalar area and the endopiriform nucleus. In summary, our three different statistical analyses are converging to show that several structures show a higher density of cFos but not of tdT neurons in the NW condition compared to Mod and Sol.

**Supp. Table 3.**
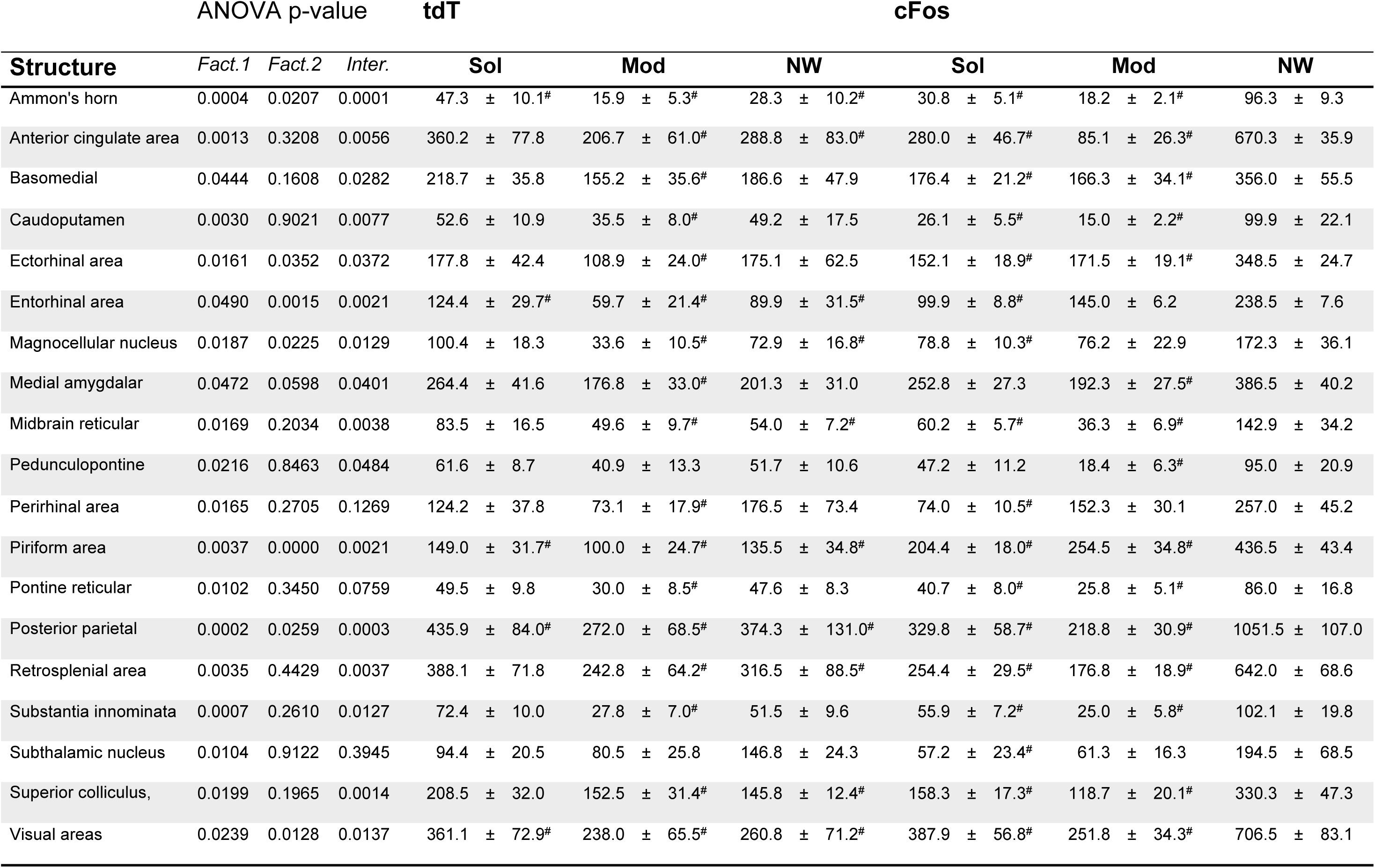
Structures increased in NW cFos condition. Columns show from the left to the right, the two way ANOVA p-values (Factor 1, treatment, Factor 2, marker and interaction between the two), the mean values of tdT and cFos cell densities per mm2 and the Standard error of the mean (SEM) for each condition and structure. The stars show p<0.05 (Tukey-Kramer post-hoc pairwise comparison) within each marker. (#) *vs* NW-cFos.

| Structure | ANOVA p-value |  |  | tdT |  |  | cFos |  |  |
| --- | --- | --- | --- | --- | --- | --- | --- | --- | --- |
|  | Fact.1 | Fact.2 | Inter. | Sol | Mod | NW | Sol | Mod | NW |
| Ammon's horn | 0.0004 | 0.0207 | 0.0001 | 47.3 ± 10.1<sup>#</sup> | 15.9 ± 5.3<sup>#</sup> | 28.3 ± 10.2<sup>#</sup> | 30.8 ± 5.1<sup>#</sup> | 18.2 ± 2.1<sup>#</sup> | 96.3 ± 9.3 |
| Anterior cingulate area | 0.0013 | 0.3208 | 0.0056 | 360.2 ± 77.8 | 206.7 ± 61.0<sup>#</sup> | 288.8 ± 83.0<sup>#</sup> | 280.0 ± 46.7<sup>#</sup> | 85.1 ± 26.3<sup>#</sup> | 670.3 ± 35.9 |
| Basomedial | 0.0444 | 0.1608 | 0.0282 | 218.7 ± 35.8 | 155.2 ± 35.6<sup>#</sup> | 186.6 ± 47.9 | 176.4 ± 21.2<sup>#</sup> | 166.3 ± 34.1<sup>#</sup> | 356.0 ± 55.5 |
| Caudoputamen | 0.0030 | 0.9021 | 0.0077 | 52.6 ± 10.9 | 35.5 ± 8.0<sup>#</sup> | 49.2 ± 17.5 | 26.1 ± 5.5<sup>#</sup> | 15.0 ± 2.2<sup>#</sup> | 99.9 ± 22.1 |
| Ectorhinal area | 0.0161 | 0.0352 | 0.0372 | 177.8 ± 42.4 | 108.9 ± 24.0<sup>#</sup> | 175.1 ± 62.5 | 152.1 ± 18.9<sup>#</sup> | 171.5 ± 19.1<sup>#</sup> | 348.5 ± 24.7 |
| Entorhinal area | 0.0490 | 0.0015 | 0.0021 | 124.4 ± 29.7<sup>#</sup> | 59.7 ± 21.4<sup>#</sup> | 89.9 ± 31.5<sup>#</sup> | 99.9 ± 8.8<sup>#</sup> | 145.0 ± 6.2 | 238.5 ± 7.6 |
| Magnocellular nucleus | 0.0187 | 0.0225 | 0.0129 | 100.4 ± 18.3 | 33.6 ± 10.5<sup>#</sup> | 72.9 ± 16.8<sup>#</sup> | 78.8 ± 10.3<sup>#</sup> | 76.2 ± 22.9 | 172.3 ± 36.1 |
| Medial amygdalar | 0.0472 | 0.0598 | 0.0401 | 264.4 ± 41.6 | 176.8 ± 33.0<sup>#</sup> | 201.3 ± 31.0 | 252.8 ± 27.3 | 192.3 ± 27.5<sup>#</sup> | 386.5 ± 40.2 |
| Midbrain reticular | 0.0169 | 0.2034 | 0.0038 | 83.5 ± 16.5 | 49.6 ± 9.7<sup>#</sup> | 54.0 ± 7.2<sup>#</sup> | 60.2 ± 5.7<sup>#</sup> | 36.3 ± 6.9<sup>#</sup> | 142.9 ± 34.2 |
| Pedunculopontine | 0.0216 | 0.8463 | 0.0484 | 61.6 ± 8.7 | 40.9 ± 13.3 | 51.7 ± 10.6 | 47.2 ± 11.2 | 18.4 ± 6.3<sup>#</sup> | 95.0 ± 20.9 |
| Perirhinal area | 0.0165 | 0.2705 | 0.1269 | 124.2 ± 37.8 | 73.1 ± 17.9<sup>#</sup> | 176.5 ± 73.4 | 74.0 ± 10.5<sup>#</sup> | 152.3 ± 30.1 | 257.0 ± 45.2 |
| Piriform area | 0.0037 | 0.0000 | 0.0021 | 149.0 ± 31.7<sup>#</sup> | 100.0 ± 24.7<sup>#</sup> | 135.5 ± 34.8<sup>#</sup> | 204.4 ± 18.0<sup>#</sup> | 254.5 ± 34.8<sup>#</sup> | 436.5 ± 43.4 |
| Pontine reticular | 0.0102 | 0.3450 | 0.0759 | 49.5 ± 9.8 | 30.0 ± 8.5<sup>#</sup> | 47.6 ± 8.3 | 40.7 ± 8.0<sup>#</sup> | 25.8 ± 5.1<sup>#</sup> | 86.0 ± 16.8 |
| Posterior parietal | 0.0002 | 0.0259 | 0.0003 | 435.9 ± 84.0<sup>#</sup> | 272.0 ± 68.5<sup>#</sup> | 374.3 ± 131.0<sup>#</sup> | 329.8 ± 58.7<sup>#</sup> | 218.8 ± 30.9<sup>#</sup> | 1051.5 ± 107.0 |
| Retrosplenial area | 0.0035 | 0.4429 | 0.0037 | 388.1 ± 71.8 | 242.8 ± 64.2<sup>#</sup> | 316.5 ± 88.5<sup>#</sup> | 254.4 ± 29.5<sup>#</sup> | 176.8 ± 18.9<sup>#</sup> | 642.0 ± 68.6 |
| Substantia innominata | 0.0007 | 0.2610 | 0.0127 | 72.4 ± 10.0 | 27.8 ± 7.0<sup>#</sup> | 51.5 ± 9.6 | 55.9 ± 7.2<sup>#</sup> | 25.0 ± 5.8<sup>#</sup> | 102.1 ± 19.8 |
| Subthalamic nucleus | 0.0104 | 0.9122 | 0.3945 | 94.4 ± 20.5 | 80.5 ± 25.8 | 146.8 ± 24.3 | 57.2 ± 23.4<sup>#</sup> | 61.3 ± 16.3 | 194.5 ± 68.5 |
| Superior colliculus, | 0.0199 | 0.1965 | 0.0014 | 208.5 ± 32.0 | 152.5 ± 31.4<sup>#</sup> | 145.8 ± 12.4<sup>#</sup> | 158.3 ± 17.3<sup>#</sup> | 118.7 ± 20.1<sup>#</sup> | 330.3 ± 47.3 |
| Visual areas | 0.0239 | 0.0128 | 0.0137 | 361.1 ± 72.9<sup>#</sup> | 238.0 ± 65.5<sup>#</sup> | 260.8 ± 71.2<sup>#</sup> | 387.9 ± 56.8<sup>#</sup> | 251.8 ± 34.3<sup>#</sup> | 706.5 ± 83.1 |

### Differential activation of catecholaminergic and hypocretin/orexin neurons in Sol, Mod or NW conditions

To determine whether classical wake-promoting populations of neurons were differentially activated by Sol, Mod and NW, triple labeling of tdT, cFos and TH or orexin (Orx) were made. Then, the number of double and triple labeled neurons was determined in all mice and groups. To statistically compare the number of double-labeled cells across groups and mice in each structure, the number of double-labeled neurons was divided by the total number of TH or Orx and then a two-way ANOVA was performed followed by a post-hoc comparison (Tukey’s honestly significant difference procedure).

For the catecholaminergic neurons, different levels of activation were observed among the different cell groups. The hypothalamic A11-A13 dopaminergic (DA) cell groups located in the zona incerta (ZI) consistently showed less than 4% of TH neurons co-expressing tdT or cFos across all conditions, with no significant differences between groups (**Table 2**). Similarly, the ventral tegmental area (VTA, A10 dopaminergic (DA) group, **Fig 8**) and the substantia nigra (SN, A9 DA group), both containing a very large number of TH-labeled neurons, showed very few double-labeled neurons. Indeed, less than 2% of the TH+ neurons in these nuclei expressed tdT or cFos, irrespective of the experimental condition (**Table 2**).

**Fig 8.**
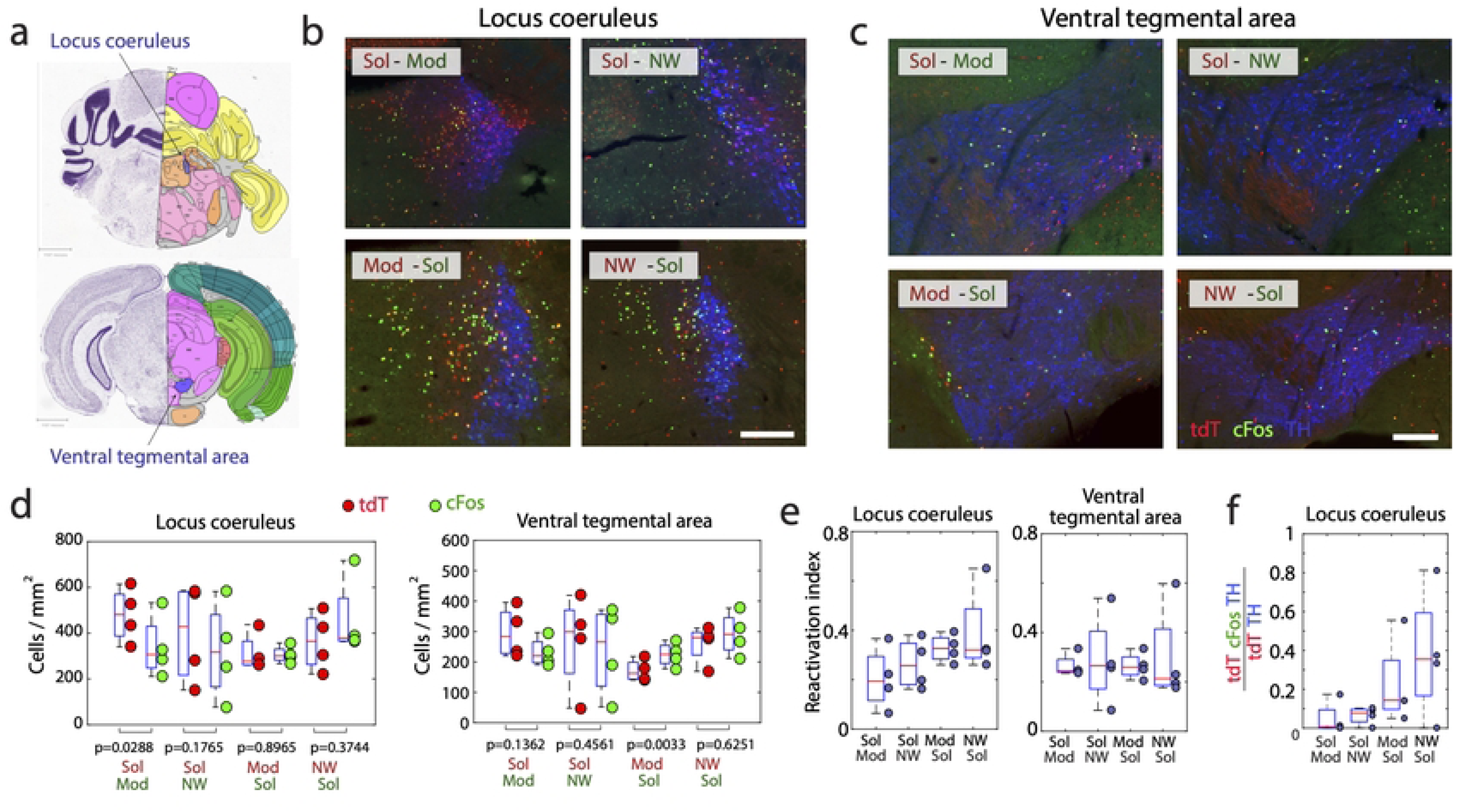
Activation of the LC and VTA catecholaminergic groups during wakefulness. (**a**) Drawings showing the localization of the two structures. (**b-c**) Photomicrographs illustrating the distribution of tdT (red), cFos (green), TH (blue), tdT and TH double-labeled neurons (purple) and cFos and TH double labeled neurons (white nuclei) in one representative mouse per experimental group in the LC and VTA. (**b**) In the LC, the number of TH and tdT or cFos double-labeled neurons is low in all conditions although it is significantly higher in the Sol condition. (**c**) A very low number of TH double-labeled neurons are visible in the VTA in all conditions. (**d**) Boxplots showing that the density (cell/mm^3^) of tdT+ (red) and cFos+(green) neurons in the LC and VTA is not different among the four groups (Sol-Mod, Sol-NW, Mod-Sol, NW-Sol, n = 4 per group). Significance two-way ANOVA followed by a post-hoc test when comparing tdT and cFos for each group. **(e)** The Reactivation index (tdTomato+/cFos+ neurons over total tdTomato+ neurons) for the two structures is also not different across the four groups. **(f)** Percentage of TH neurons double-labeled with tdT or cFos in the LC. Note in all conditions the moderate activation in the LC and the very low level of activation of the TH neurons in the VTA (not shown since more than half of the mice had no tdT-cFos-TH+ cells). Significance one-way Anova and Post-hoc test. p < 0.05*. Boxes, median; dots, individual mouse values; boxes, first and last quartiles; whiskers, minimum and maximum values excluding outliers. Scale b: 300 μm; scale c: 200 μm.

**Table 2.** Catecholaminergic neurons. The table shows the percentage of TH neurons double labeled ([tdT-TH+]/TH+ or [Fos-TH+]/TH+) after treatment with Sol, Mod or non-pharmacological wakefulness (NW). Data shown as mean ± SEM. Sol groups N=8, Mod and NW groups, N=4. The two-way ANOVA p-values (factor 1: experimental treatment; factor 2: marker, and interaction between the two factors) are shown. (*) *vs* NW, (**) *vs* Sol, and (+) *vs* Sol-tdT groups.

| Structure | ANOVA p-value |  |  | tdT |  |  | cFos |  |  |
| --- | --- | --- | --- | --- | --- | --- | --- | --- | --- |
|  | Fact.1 | Fact.2 | Inter. | Sol | Mod | NW | Sol | Mod | NW |
| Dorsal nucleus raphe | 0.250 | **0.043** | 0.072 | 20.11 ± 3.64 | 13.05 ± 2.83 | 11.56 ± 2.78 | 7.75 ± 2.67 + | 3.74 ± 1.43 + | 15.03 ± 3.80 |
| Lateral reticular nucleus | **0.001** | 0.339 | 0.232 | 54.68 ± 5.46 | 30.28 ± 3.35 | 44.71 ± 13.26 | 61.57 ± 7.03 | 23.32 ± 4.13 ** | 26.11 ± 7.90** |
| Locus ceruleus | **0.012** | **0.024** | 0.150 | 18.91 ± 3.30 | 6.37 ± 1.73 | 6.15 ± 1.71 | 6.34 ± 3.04 + | 1.50 ± 1.20 + | 4.98 ± 1.68+ |
| N. of the solitary tract | **0.004** | 0.191 | 0.135 | 27.10 ± 3.84 | 15.61 ± 3.27 | 26.06 ± 10.71 | 32.86 ± 4.46 | 5.52 ± 2.40 ** | 12.00 ± 5.68** |
| Substantia nigra, *pars c.* | 0.441 | 0.222 | 0.665 | 0.22 ± 0.14 | 0.16 ± 0.16 | 0.00 ± 0.00 | 0.04 ± 0.04 | 0.00 ± 0.00 | 0.00 ± 0.00 |
| Ventral tegmental area | 0.143 | 0.786 | 0.795 | 0.93 ± 0.30 | 0.08 ± 0.08 | 0.91 ± 0.49 | 1.44 ± 0.74 | 0.10 ± 0.10 | 0.75 ± 0.25 |
| Zona incerta | 0.433 | 0.946 | 0.831 | 2.50 ± 1.18 | 1.82 ± 0.79 | 1.84 ± 0.92 | 3.18 ± 0.84 | 1.96 ± 1.24 | 1.21 ± 0.79 |

In the dorsal raphe nucleus (DRN), the median percentage of TH+ neurons co-expressing tdT or cFos was higher than in the above dopaminergic structures (∼12%), but not statistically different between conditions (**Table 2**). The median percentage of locus coeruleus TH+ neurons (LC, A6 noradrenergic, NA group, **Fig 8**) co-expressing tdT or cFos was also only 12 % and 5%, respectively. Nevertheless, LC neurons were differentially affected by the experimental treatment (p=0.012, factor 1, 2-way ANOVA, **Table 2**) with a significantly higher percentage of TH neurons double labeled in Sol-treated animals compared to the two other conditions (19% Sol-tdT+/TH+ and 6% Sol-cFos+/TH+, **Table 2**). However, the Tukey post-hoc test did not show a statistical difference between conditions.

In the nucleus of the solitary tract (NTS, containing the A2 NA group, **Fig 7**), the median percentage of TH+ neurons co-expressing tdT or cFos in all mice was of 20%, a higher percentage than for the above catecholaminergic groups. Two-way ANOVA revealed a treatment effect (p=0.004, factor 1, **Table 2**) with Sol-treated groups showing a higher percentage of TH neurons double-stained compared to the two other groups (27% Sol-tdT+/TH+ and 33% Sol-cFos+/TH+). It did reach statistical difference when comparing the percentage of cFos+/TH+ neurons between the three groups but not when comparing the percentage of tdT+/TH+ neurons (**Table 2**). The paired t test also showed a statistically higher number of tdT neurons compared to cFos in the Sol-NW group.

In the lateral reticular nucleus (LRN, A1 NA group, **Fig 7**), the median percentage of TH+ neurons co-expressing tdT or cFos was over 50% in the Sol condition and between 20-40% in the NW and Mod conditions (**Fig 7e**, **Table 2**). Two-way ANOVA showed that Sol-treated mice have a higher percentage of TH+ neurons expressing tdT or cFos than Mod and NW groups (p=0.001, factor 1, 2-way ANOVA, **Table 2**). It did reach statistical difference using Tukey post-hoc test for the cFos double-labeled neurons but not the Tdt ones. The paired t test showed a statistically higher number of tdT neurons compared to cFos in the Sol-Mod group and conversely in the NW-Sol group **(Fig 7i)**.

For the hypocretin/orexin (hcrt-orx) cell group located in the lateral hypothalamic area (LHA, (**Fig 10**), the median percentage of activated hcrt-orx neurons (i.e., labeled with tdT or cFos) across all mice was of 38% **(Fig 10e)**, indicating a high level of activation during the Mod, Sol and NW conditions with no statistical difference between conditions. Overall, the median percentage of tdT/orx neurons co-expressing cFos was 39.5% (**Fig 10h**), while that of cFos/orx neurons co-expressing tdT was 70.2% (**Fig 10i**). Orx neurons constituted 36% and 21% of the tdT and cFos population in the LHA, respectively.

### Structures strongly activated in all conditions

Our final aim was to identify populations of neurons strongly activated in the three different conditions namely Sol, Mod and NW to identify candidate structures implicated in inducing wakefulness in all conditions. For that, we selected structures with high (>0.3) averaged RI in all of the four groups of mice (**Table 3**). The structures with a RI superior to 0.3 in the four groups of mice were mostly composed of hypothalamic nuclei including the parasubthalamic nucleus (**Fig 9**), posterior hypothalamic nucleus, supramammillary nucleus and the ventromedial and dorsomedial nuclei of the hypothalamus. Additionally, three brainstem nuclei namely the laterodorsal tegmental nucleus, the lateral reticular nucleus (**Fig 7c,f,i**) and the tegmental reticular nucleus which is known to be involved in motor control were selected. One thalamic nucleus implicated in the circadian system, the geniculate group-ventral thalamus, and two nuclei from the sensory system, the inferior and the superior colliculus sensory related, were also strongly reactivated in all conditions (not illustrated). The lateral hypothalamic area and the paraventricular nucleus of the thalamus could also be added to these structures since they displayed a high density of cells in all three conditions and their reactivation index was above or close to 0.3 in all conditions **(Fig 9d,e, 10d,f)**. In summary, our results reveal structures not previously implicated in wake control excepting the laterodorsal tegmental nucleus and the Orx neurons and therefore point out that the previously identified structures might not be the most important to generate the state.

**Table 3.** Listing of the structures showing a high reactivation index in all groups. Structures with averaged reactivation index above 0.3 in all groups and with a p > 0.05 in one-way ANOVA (d.f.=3,12), excepting the Lateral Reticular Nucleus showing just a value just above (0.0051). The numbers in each group show the mean ± SEM.

| Structure | ANOVA p-value |  | Reactivation index |  |  |  |
| --- | --- | --- | --- | --- | --- | --- |
|  | F | p-val. | Sol-Mod | Sol-NW | Mod-Sol | NW-Sol |
| Dorsomedial nucleus of | 1.93 | 0.2268 | 0.36 ± 0.03 | 0.32 ± 0.00 | 0.50 ± 0.05 | 0.48 ± 0.04 |
| Geniculate group. ventral | 0.37 | 0.7735 | 0.44 ± 0.03 | 0.56 ± 0.06 | 0.50 ± 0.11 | 0.53 ± 0.09 |
| Inferior colliculus | 2.06 | 0.1598 | 0.34 ± 0.03 | 0.49 ± 0.05 | 0.43 ± 0.04 | 0.48 ± 0.05 |
| Lateral reticular nucleus | 7.54 | **0.0051** | 0.39 ± 0.03 | 0.33 ± 0.04 | 0.54 ± 0.02 | 0.37 ± 0.02 |
| Laterodorsal tegmental | 3.22 | 0.0612 | 0.38 ± 0.03 | 0.45 ± 0.05 | 0.43 ± 0.03 | 0.56 ± 0.03 |
| Parasubthalamic nucleus | 1.04 | 0.4124 | 0.42 ± 0.07 | 0.54 ± 0.02 | 0.57 ± 0.05 | 0.47 ± 0.06 |
| Posterior hypothalamic | 1.52 | 0.2602 | 0.31 ± 0.05 | 0.52 ± 0.05 | 0.44 ± 0.05 | 0.40 ± 0.08 |
| Superior colliculus. | 0.50 | 0.6900 | 0.37 ± 0.06 | 0.44 ± 0.00 | 0.34 ± 0.08 | 0.39 ± 0.03 |
| Supramammillary nucleus | 0.38 | 0.7720 | 0.34 ± 0.06 | 0.48 ± 0.00 | 0.47 ± 0.09 | 0.46 ± 0.04 |
| Tegmental reticular | 0.97 | 0.4432 | 0.44 ± 0.08 | 0.54 ± 0.08 | 0.49 ± 0.10 | 0.31 ± 0.03 |
| Ventromedial | 0.96 | 0.4447 | 0.34 ± 0.04 | 0.53 ± 0.10 | 0.49 ± 0.06 | 0.44 ± 0.08 |

**Fig 9.**
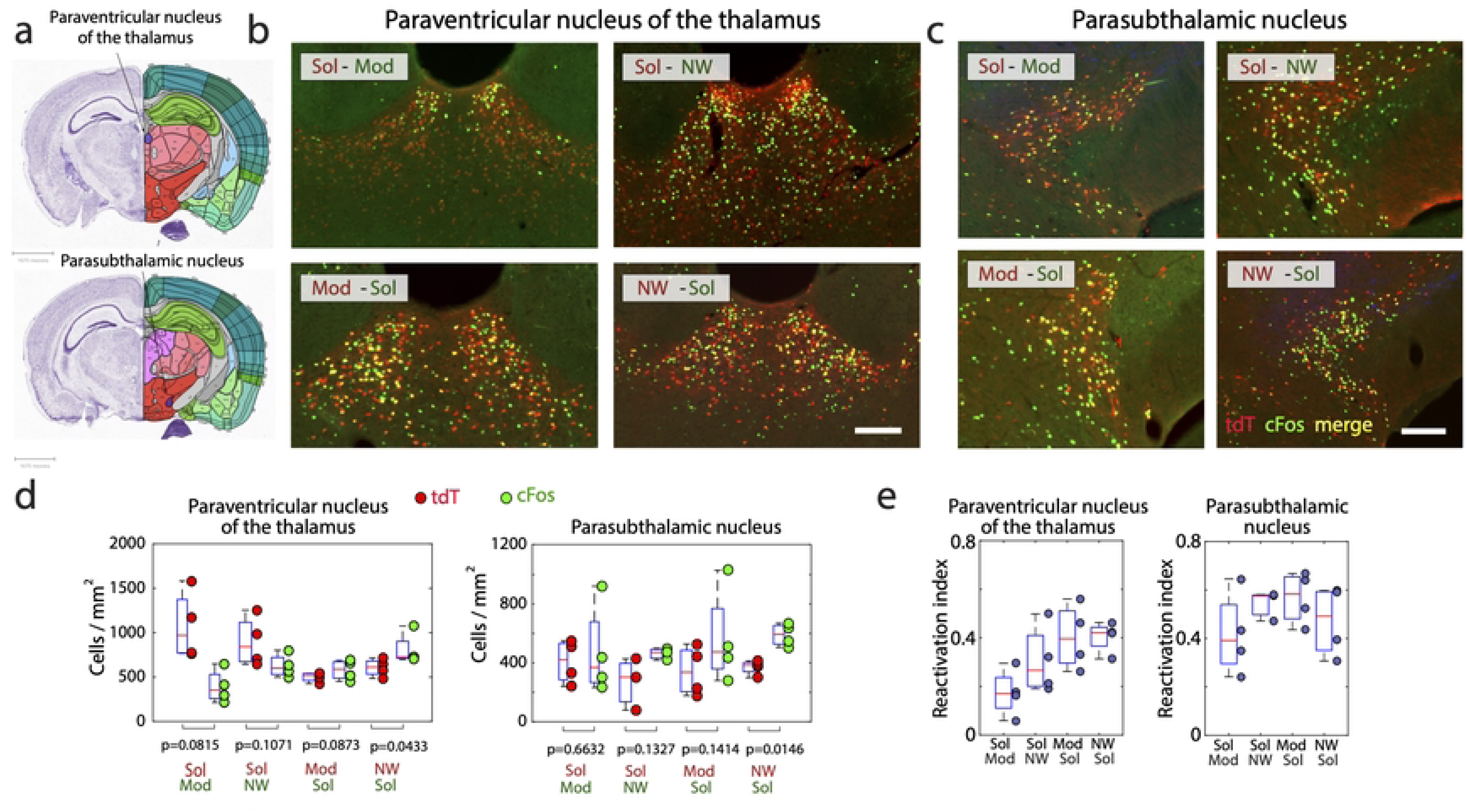
The lateral hypothalamic area and the Hcrt/Orx neurons are strongly activated during all types of wakefulness. (**a**) Drawings showing the localization of the LHA. (**b-c**) Low power and high power magnification (b) of photomicrographs illustrating the distribution of tdT (red), cFos (green), Orx (blue), tdT and Orx double-labeled neurons (purple), and cFos and Orx double labeled neurons (white nuclei) in one representative mouse per experimental group. Note that in all groups of mice, many cFos, tdT and double-labeled and triple-labeled neurons are labeled in the LHA. (**d**) Boxplots showing the density (cell/mm^3^) of tdT+ (red) and cFos+ (green) neurons in the lateral hypothalamic area in each of the four groups (Sol-Mod, Sol-NW, Mod-Sol, NW-Sol, n=4 per group). Significance two-way anova followed by a post-hoc test when comparing tdT and cFos for each group. **(e)** Percentage of Orx neurons double-labeled with tdT or cFos in the LHA. **(f)** Reactivation index (tdTomato+/cFos+ neurons over total tdTomato+ neurons) in the LHA. Note the high level of reactivation in all conditions except for the Sol-Mod condition showing a non-significant lower RI. **(g)** Percentage of Orx neurons triple labeled **(h)** Percentage of tdT-Orx double-labeled which are cFos+ **(i)** Percentage of cFos-Orx neurons which are triple labeled. Significance one-way Anova and Post-hoc test. p < 0.05*. Boxes, median; dots, individual mouse values; boxes, first and last quartiles; whiskers, minimum and maximum values excluding outliers. Scale 100 μm.

**Fig 10.**
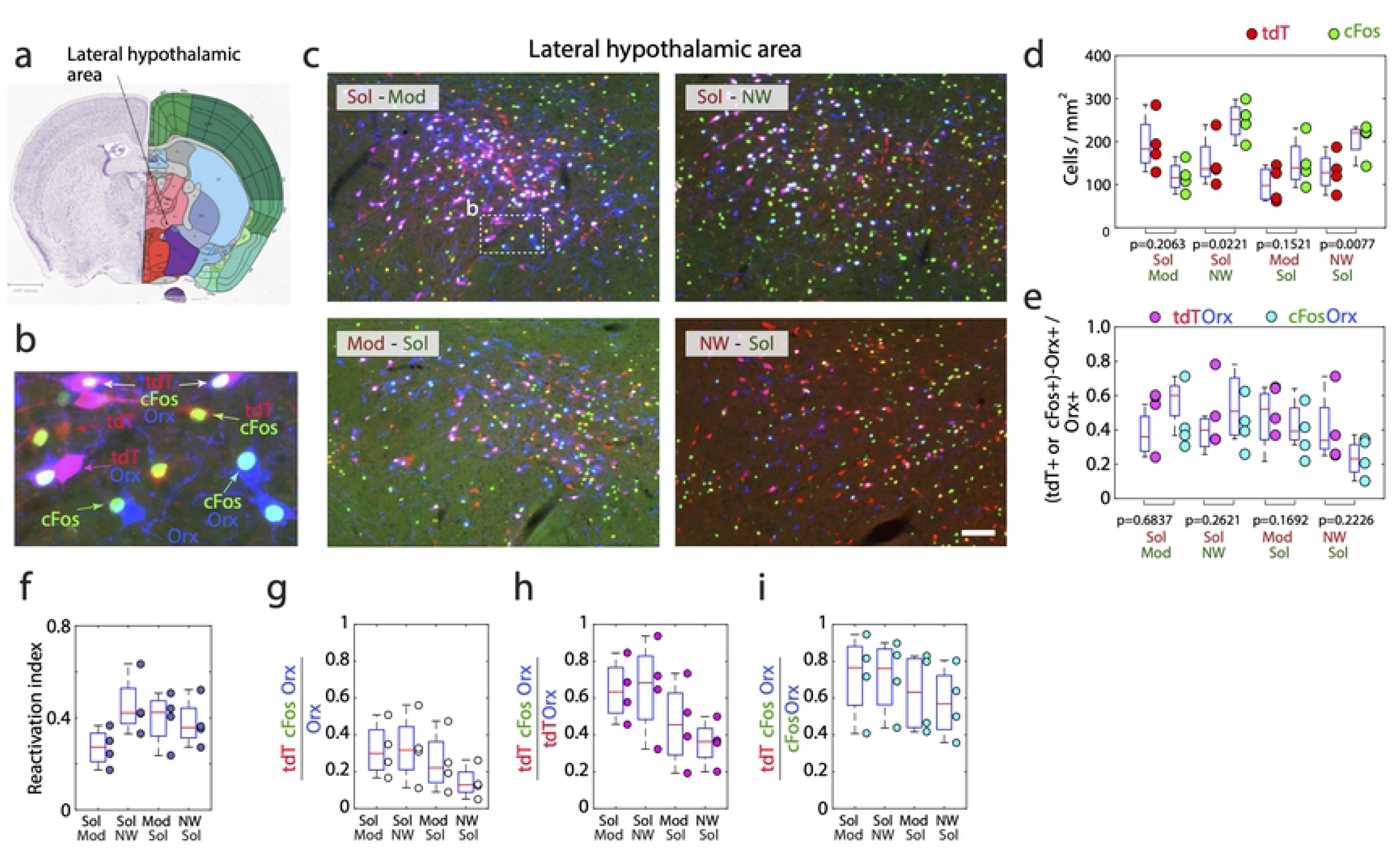
Strong activation of the PVT during Sol and of the PSTN during all conditions. (**a**) Drawings showing the localization of the two structures illustrated. (**b-c**) Photomicrographs illustrating the distribution of tdT (red cytoplasm), cFos (green nuclei) and double-labeled neurons (yellow cytoplasm) in one representative mouse per experimental group in the paraventricular thalamic nucleus (PVT) and the parasubthalamic nucleus (PSTN). Note that for the PSTN, many cFos, tdT and double-labeled neurons are labeled in all conditions. (**d**) Boxplots showing the density (cell/mm^3^) of tdT+ (red) and cFos+(green) neurons in each of the four groups (Sol-Mod, Sol-NW, Mod-Sol, NW-Sol, n = 4 per group). Significance two-way anova followed by a post-hoc test when comparing tdT and cFos for each group. **(e)** High reactivation index (tdT+/cFos+ neurons over total tdT+ neurons) for the two structures except for the Sol-Mod and the Mod-Sol conditions for the PVT. Significance one-way Anova and Post-hoc test. p < 0.05*. Boxes, median; dots, individual mouse values; boxes, first and last quartiles; whiskers, minimum and maximum values excluding outliers. Scale 200 μm.

## Discussion

Our results indicate that a set of subcortical structures are significantly more activated during wakefulness induced by Solriamfetol than by Modafinil and non-pharmacological wakefulness suggesting that Sol induced a differential brain activation than the two other conditions. Nevertheless, some subcortical structures were also strongly activated during all three conditions suggesting also some common mechanisms for wake induction by the three treatments. We discuss below the significance of these results in detail including methodological considerations and physiological implications.

### Methodological considerations

To our knowledge, our study is the first to use the TRAP mice to compare the neuronal activation occurring during wakefulness induced by two different drugs versus behavioral stimulation. More generally, our study is the first to extensively compare the distribution of the reporter gene staining obtained in the TRAP mice with endogenous cFos expression. Our paradigm assumes that tdT labeling induced by the removal of the stop codon on the inserted tdTomato gene by the CreERT2 transgene under the cFos promoter after 4-OHT injection is comparable to natural cFos expression. To rule out the possibility of potential differences between the two methods, we inverted the conditions in four groups of four mice (Mod-Sol versus Sol-Mod and NW-Sol versus Sol-NW). By this means, we were able to rule out wrong conclusions since we did compare the number of tdT and cFos+ cells inside each group but also the number of cFos+ and tdT+ for each condition independently. Moreover, we also did compare the percentage of double-labeled neurons between groups using the reactivation index. First, as shown in Fig 2, the correlation between the number of tdT and cFos labeled neurons in each structure and for all mice was very high indicating that the two methods are close to each other. Second, we decided to highlight only the results showing a reproducible modification across markers, groups of mice and conditions. There were only two instances for which tdT and cFos staining did not match. First, there were statistically more cFos than tdT labeled neurons in several structures mostly cortical in the NW-Sol group but there was not more tdT than cFos neurons conversely in the Sol-NW group. This finding suggests that the sensitivity of the tdT staining is lower than that of cFos, in particular in the cortex and only for the NW condition. It can be hypothesized that the induction of tdT expression might be lower specifically in the NW condition. One possibility is that the efficiency of the TRAP method is lower particularly in the cortex and for specific conditions such as NW. On the other hand, six thalamic structures showed an increased number of tdT cells in the Sol condition compared to the NW and Mod conditions. Surprisingly, these structures did not show an increased number of cFos-positive neurons compared to tdT in the NW-Sol or the Mod-Sol groups. Such discrepancy suggests a higher sensitivity of tdT staining in these thalamic nuclei. It indicates that although the cFos and the TRAP methods are quite similar, control experiments should be done to confirm that they give the same results for the structures modified by the conditions under study.

In summary, while there are some discrepancies between the two methods, overall, they produce similar and comparable results in most structures, in particular in those showing a differential activation with Sol compared to Mod and NW or on the contrary showing a high reactivation index in all groups of mice.

### Comparison with previous studies

There is only one previous study in which they compared the distribution of cFos induced by solriamfetol, modafinil and non-pharmacological wakefulness. They reported that both drugs shared many areas of similar activation such as the lateral septum, the septohippocampal nucleus, the dorsal tenia tecta, the semilunar nucleus, the island of calleja, the olfactory tubercle, and the arcuate nucleus (Hasan et al., 2009). The subiculum, ventral tenia tecta, the nucleus of solitary tract gelatinous, dorsal endopiriform, and suprachiasmatic nucleus were activated by solriamfetol, specifically. It is difficult to compare these results with the present ones since the doses used were much higher (150 mg/kg vs 32 for Mod and 60mg/kg for Sol) and the analysis was not detailed and lacked statistical analysis (Hasan et al., 2009). In addition, cFos labeling was analyzed in four previous publications after modafinil injection but again the dosage was different than ours, the mapping was compared with saline injection (i.e an undetermined state) and the analysis was not complete like in our experiments (Cruces-Solis et al., 2020; T. M. Engber et al., 1998; T.M. Engber et al., 1998; Lin et al., 1996). There were also a number of studies looking at cFos expression after periods of wakefulness induced by different methods in cats, rats and mice but the mapping was either not complete or not detailed enough to compare the data with the present study (Cirelli & Tononi, 2000; Ledoux et al., 1996; Terao et al., 2003). Finally, our study is the first in which a direct comparison is made in the same mouse between drugs and non-pharmacological wakefulness.

### Physiological significance

Our results show that the lateral septum (LS), bed nucleus of the stria terminalis (BNST), medial and median preoptic nucleus (MPO), central amygdalar nucleus (CEA), paraventricular thalamic (PVT) and hypothalamic nuclei (PVN), periventricular region, supraoptic nucleus (SON), tuberal nucleus, lateral parabrachial nucleus (LPB), nucleus of the solitary tract (NTS), area postrema and lateral reticular nucleus are significantly more activated by solriamfetol than modafinil and non-pharmacological wakefulness. Such result is puzzling since it is traditionally thought that wake is induced by classical structures such as the hypocretin/orexin and histaminergic neurons, dorsal raphe serotonergic neurons, telencephalic and pontine cholinergic neurons and the noradrenergic neurons of the locus coeruleus (Sulaman et al., 2023). However, most of the structures reported in the present study have been shown to play a role in wake mostly in very recent publications (see below). Further, there is ample evidence that these structures are interconnected (see below). Indeed, it has been shown using calcium imaging and cFos staining, that the lateral septum contains GABAergic neurons active during wake and that their optogenetic or chemogenetic activation induced wake while conversely their inhibition induced NREM (Wang et al., 2021). Further, it has been shown that wake is also induced by lateral septum projection to the GABAergic neurons of the VTA (Kodani et al., 2017). Optogenetic or chemogenetic excitation of GABAergic neurons of the BNST induces wake and such effect is attenuated by pretreatment with orexin antagonist (Kodani et al., 2017). The authors showed that the BNST strongly projects to other structures highly activated in the present paper such as the CEA, PVN, SON, LHA, NTS and LPB.

Using cFos and unit recordings, it was previously shown that PVT neurons are more active during wake than sleep (REM and moreover NREM sleep, (Ren et al., 2018). Optogenetic activation of PVT neurons and their projections to the nucleus accumbens induces wake whereas their lesion and optogenetic or chemogenetic inactivation reduces it, while increasing NREM during the dark period (Ren et al., 2018). Also, chemogenetic inactivation of the hypocretin/orexin (Orx) projection to the PVT decreases wake and increases NREM. Nevertheless, wake induced by stimulating Orx neurons was not abolished by PVT chemogenetic inactivation, indicating that other pathways are involved (Ren et al., 2018). On the other hand, calcium imaging showed that PVT neurons projecting to the CEA are specifically activated during wake and that their optogenetic activation induces wakefulness while their inhibition increases NREM sleep (Zhao et al., 2022). Such pathway may mediate stress response during wake, since their optogenetic inhibition also alleviated the hormonal and behavioral responses to acute stress (Zhao et al., 2022).

The paraventricular hypothalamic nucleus (PVN) was also shown using cFos or calcium imaging to contain glutamatergic (including vasopressin, oxytocin and corticotropin releasing factor (CRH) subtypes) neurons active during wake and inactive during sleep (Chen et al., 2021). Optogenetic activation of these neurons or their projections to the lateral parabrachial nucleus (LPB) and lateral septum (LS) induced wake while their chemogenetic inhibition or ablation reduced wake and increased NREM (Chen et al., 2021). It has also been shown that chemo and optogenetic activation specifically of the vasopressin subtype of PVN neurons induces wake while their inhibition decreases it. The increase in wake was also obtained by stimulating PVN terminals in the lateral hypothalamic area and was strongly decreased by pretreatment with an Orx antagonist (Islam et al., 2022). It was also shown that restraint stress causes strong cFos expression in corticotropin releasing hormone neurons (CRH) of the PVN and also of Orx neurons (Li et al., 2020). Optogenetic stimulation of the CRH neurons of the PVN projecting to the lateral hypothalamic area leads to insomnia while inhibition of the pathway compromises restraint stress-induced insomnia (Li et al., 2020). Besides, chemogenetic and optogenetic activation of glutamatergic and calcitonin gene-related peptide (CGRP) neurons of the lateral parabrachial nucleus was also shown to induce W (Kaur et al., 2017). Conversely, optogenetic inactivation of these neurons or of their projections to the BNST, CEA and LHA increased the latency to arousal induced by hypercapnia (Kaur et al., 2017). In addition, our study identified additional structures never implicated in wake control before such as the medial and median preoptic nuclei, periventricular region, supraoptic nucleus (SON), tuberal nucleus, nucleus of the solitary tract (NTS), area postrema and lateral reticular nucleus. Interestingly, these structures have known direct neuroanatomical links to those mentioned above and therefore seem to be part of the network strongly activated by Sol.

In summary, recent published data strongly fit with our results indicating that the structures activated by Sol revealed in the present study form a network likely inducing wake **(Fig 11)**. Importantly, these structures are significantly less activated during NW and moreover Mod indicating that Sol has a different mode of action than Mod. This is surprising since both drugs are known to be norepinephrine and dopamine reuptake blockers. However, it has recently been shown that Sol but not Mod has additional agonist activity at the trace amine associated receptor 1 (TAAR1) (Gursahani et al., 2022). Besides, it has been shown that a TAAR1 agonist induces W and inhibits NREM sleep (Schwartz et al., 2017). In addition, increased cFos staining has been reported after injection of two different TAAR1 agonists in several structures specifically activated by Sol in our study namely the bed nucleus of the stria terminalis, central amygdala, lateral parabrachial nucleus, nucleus of the solitary tract, area postrema, parasubthalamic nucleus (PSTN) and lateral hypothalamic area (Dedic et al., 2024). Altogether, these data strongly suggest that part of the wake inducing effect of Sol is mediated through activation of a set of structures by its TAAR1 receptors agonist activity.

**Fig 11.**
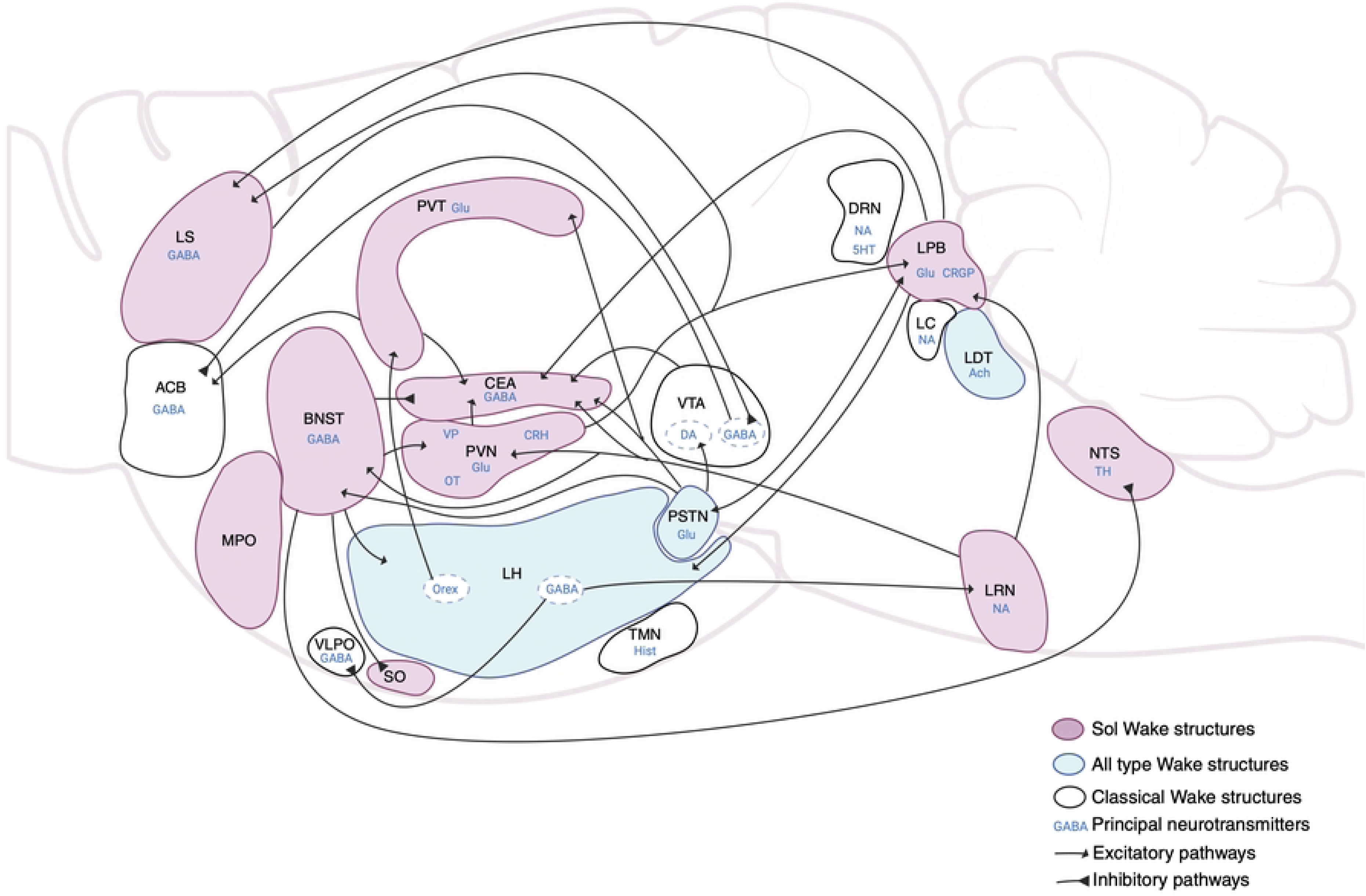
The neuronal network of the wakefulness system activated by Sol, Mod and NW. Drawing showing many interconnected structures implicated in wakefulness, after Sol, and Mod injections and NW. The pink structures are significatively more activated by Sol than Mod and NW. Sol strongly activates a subset of structures less activated in non-pharmacological wakefulness (NW) and only weekly activated by Mod. The blue structures are activated by Sol, Mod and NW. The white ones are known to be implicated in the wake system, for the DRN, LC and TMN, their role is minor. Different neurotransmitters are involved in this system inhibiting or exciting wake structures, like gamma-aminobutyric (GABA), glutamate (Glu), noradrenalin (NA), Histamine (Hist), acetylcholine (Ach), Orexin (Orx), dopamine (DA), tyrosine hydroxylase (TH), serotonin (5HT), oxytocin (OT), vasopressin (VP), corticotropin-releasing hormone (CRH), and calcitonin gene-related peptide (CRGP). ACB: accumbens nucleus, BNST: bed nucleus of the stria terminalis, CEA: central amygdala, DRN: dorsal raphe nucleus, LC: locus coeruleus, LDT: laterodorsal tegmental nucleus, LH: lateral hypothalamus, LPB: lateral parabrachial nucleus, LRN: lateral reticular nucleus, LS: lateral septum, MPO: medial preoptic nucleus, NTS: nucleus of the solitary tract, PSTN: parasubthalamic nucleus, PVN: paraventricular nucleus of the hypothalamus, PVT: paraventricular nucleus of the thalamus, SO: supraoptic nucleus, TMN: tuberomammillary nucleus, VLPO: ventrolateral preoptic area, VTA: ventral tegmental area.

In addition, the parasubthalamic nucleus, lateral hypothalamic area, laterodorsal tegmental nucleus, posterior hypothalamic nucleus, supramammillary nucleus, dorsomedial and ventromedial nuclei of the hypothalamus, geniculate group of the ventral thalamus, superior colliculus, sensory related, tegmental reticular nucleus and the lateral reticular nucleus showed a high reactivation index in all groups of mice indicating that they are strongly activated during wake induced by Sol, Mod and NW. The geniculate group of the ventral thalamus, the superior colliculus, sensory related nuclei are known to be implicated in circadian or sensory processing and are therefore unlikely involved in specifically inducing wake. In contrast, some of the other structures identified have been recently implicated in wake induction. Indeed, glutamatergic neurons of the parasubthalamic nucleus (PSTN) are active during W and REM sleep but not NREM (Guo et al., 2023). The optogenetic and chemogenetic activation of these neurons and their projection to the VTA and lateral parabrachial nucleus increases wakefulness and exploratory behavior while their inhibition decreases wakefulness (Guo et al., 2023). The PSTN receives an excitatory projection from parabrachial glutamatergic neurons. The PSTN also projects to the bed nucleus of the stria terminalis, central amygdala and PVT (Guo et al., 2023). The lateral hypothalamic area (LHA) also contained a large number of neurons activated by Sol, Mod and NW. We further showed in the present study that 26% of these neurons were Hcrt/Orx neurons. The role of Hcrt/Orx neurons in wake induction and maintenance is supported by a large number of studies showing using cFos and unit recordings that they are specifically active during W and that their chemogenetic and optogenetic activation induces W, while their inhibition or ablation leads to hypersomnia (Sulaman et al., 2023). In addition to the Hcrt/Orx neurons, a population of GABAergic neurons of the LHA have been shown to be W active and their optogenetic and chemogenetic activation as well as that of their projection to sleep-active structures such as the lateral reticular thalamic nucleus and the ventrolateral preoptic area induced wake (Herrera et al., 2016; Venner et al., 2016). Finally, it has also been previously shown that the laterodorsal tegmental nucleus, in particular its cholinergic neurons, plays a role in W induction (Manns et al., 2000). The chemogenetic activation and inhibition of the supramammillary nucleus was also shown to increase or decrease wakefulness, respectively (Pedersen et al., 2017). Finally, to our knowledge the posterior hypothalamic nucleus, and dorsomedial and ventromedial nuclei of the hypothalamus have not been previously involved in W and additional data is needed to determine whether they could play a role.

We did also determine whether the dopaminergic and noradrenergic systems are activated during wake induced by Mod, Sol and NW. Surprisingly, no more than 4% of the dopaminergic neurons of the hypothalamic A11 and A13 (zona incerta) and the mesencephalic A10 (VTA) and A9 (substantia nigra) groups were activated in all conditions. These results are in line with our previous cFos study in rats showing that these neurons are not activated during W (Leger et al., 2010). They are also supported by calcium imaging of DA neurons of the VTA showing their low activity during W although it was higher than during NREM sleep (Eban-Rothschild et al., 2016). Nevertheless, optogenetic activation of these neurons or their projection to the nucleus accumbens, the striatum and the central amygdala induced W while their inhibition promoted nest building, induced a decrease in wake and increase in NREM indicating that despite their low activity, they play a role in wake induction (Eban-Rothschild et al., 2016). In the dorsal raphe nucleus (DRN), only 12% of the dopaminergic neurons did express tdT or cFos in all three wake conditions suggesting that they might play a minor role in inducing the wake state. Fiber photometry reported that dopaminergic neurons of the DRN are indeed specifically active at the onset of W. Optogenetic activation of dopaminergic neurons of the DRN induced wake while their chemogenetic inactivation decreased it and increased NREM (Cho et al., 2017). Based on these and our data, it is likely that only a subset of DA neurons of the DRN are involved in wake control.

In the case of the LC noradrenergic neurons, NA A6 group), less than 12% of the neurons were activated in all conditions. In our previous study in rats, we reported that 30% of the noradrenergic LC neurons were cFos+ after wake induced by a novel environment with two rats in the arena (Leger et al., 2009). The higher number of double-labeled neurons in the previous study might be due to the stronger stimulation induced by social exposure. Optogenetic excitation of LC noradrenergic neurons induced wake while their inhibition induced a small reduction of wake (Carter et al., 2010). Further, optogenetic inhibition decreases sensory evoked awakenings (Hayat et al., 2020). Interestingly, cFos staining induced during wake was strongly decreased in the cortex but not the hypothalamus after LC lesion (Cirelli et al., 1996). Altogether, these results suggest that LC noradrenergic neurons directly activate the cortex during wake but might play a more marginal role in inducing wake by interacting with the other subcortical wake-inducing systems.

In the nucleus of the solitary tract (A2 NA group), the mean percentage of TH neurons co-expressing tdT or cFos was 30% for the Sol condition, 10% for the Mod condition and 18% for the NW condition. In the lateral reticular nucleus (A1 NA group), the median percentage of TH neurons co-expressing tdT or cFos was superior to 50% in the Sol condition and between 20-40% in the NW and Mod conditions. In our previous study in rats, around 20% of the noradrenergic neurons of A1 and A2 groups were labeled with cFos after wake (Leger et al., 2009). Although the role of A1 and A2 in inducing W is mentioned in a review on noradrenergic functions (Berridge et al., 2012), no study is devoted to such involvement. Interestingly, A1 and A2 noradrenergic neurons are strongly projecting to the BNST, CEA, PVN, SO and LPB (Woulfe et al., 1990) shown here to be statistically more activated by Sol than NW or Mod. From these results, we propose that these noradrenergic projections are playing a key role in inducing wake and are strongly recruited by Sol and to a lesser extent Mod both known to be dopaminergic and noradrenergic reuptake inhibitors (Baladi et al., 2018).

## Conclusions

In summary, our results disclose for the first time over the entire brain the neuronal systems activated by Sol, Mod and NW. In particular, we showed that Sol strongly activates a subset of structures less strongly activated in NW and only weekly activated by Mod. These results indicate that W induced by Sol is different from that induced by Mod and NW. It suggests that patients treated by Sol might be in a different neuronal state than those treated by Mod. Further, our results indicate that a large number of interconnected structures not classically involved in inducing W might play a major role in inducing the state. We created a schematic drawing showing the structures identified in the present study and the pathways likely involved (**Fig 11**). Our results therefore introduce a new concept that wake induction might be due to many more structures than previously thought and is not only induced by classical catecholaminergic, serotonergic, histaminergic and hypocretin systems. It opens the way to future studies to determine the respective roles of the newly discovered structures in wake and to study the specificity of the wake state induced by Sol.

## Material and Methods

### Animals

All experiments were conducted in accordance with the French and European Community guidelines for using animals in research and were approved by the institutional animal care and committee of the University of Lyon 1 and NEUROCAMPUS (APAFIS #31722-2021051809304729). Both male and female of double heterozygous Fos2A-iCreER;R26Ai14 (TRAP2) mice, kindly gifted by Dr. Liqun Luo from Stanford University were used. TRAP2-RED mice were generated by crossing Fos2A-iCreER/+ (TRAP2) mice to R26AI14/+ (AI14) mice [1]. In all experiments, 8-12 weeks old mice were prepared for surgery. Transgenic mice were housed individually and placed under a constant light/dark cycle (light on from 7:00 a.m. to 7:00 p.m.) and were habituated to be handled every day.

### Surgery

Mice were anesthetized with Ketamine and Xylazine (100/10 mg/kg. i.p.). Then, the top of the head was shaved, and the mice were placed in a stereotactic frame with a heating pad underneath. Two stainless screws were fixed in the parietal part (AP: -2.0 mm, ML: 1.5 mm from bregma) and one in the frontal part (AP: +2.0 mm, ML: 1.0 mm from bregma) of the skull, whereas the reference electrode for unipolar EEG recording was fixed in the occipital part (AP: -5 mm, ML: 0.0 mm from bregma). Two wire electrodes were inserted into the neck muscles for bipolar electromyogram (EMG) recordings. All leads were connected to a miniature plug (Plastics One, Roanoke, VA) that was cemented on the skull.

### Polysomnography and analysis of vigilance states

Animals were allowed to recover from surgery for 5 days in their home cage before being habituated to the recording conditions for seven days. They were then connected to a cable attached to a slip-ring commutator to allow free movement within the barrel. Unipolar EEG and bipolar EMG signals were amplified, respectively, 1:5000 V/V and 1:2000 V/V (MCP-PLUS, Alpha-Omega Engineering, Israel), digitized at 1024 Hz, and acquired using Slip Analysis v 2.9.8 software (Viewpoint, Civrieux, France). The analysis was then done as previously described (Lee et al., 2020) with slight modifications. Briefly, vigilance states of 5 s episodes were scored by a visual check of polygraphic signals (EEG and EMG recordings) according to the criteria described. Baseline sleep recordings were made during 48h starting just after the habituation. The sleep-wake cycle was then recorded for 12h before modafinil and solriamfetol injections or the exposure to the open field. The recordings were pursued for 12h in the case of 4-hydroxytamoxifen (4-OHT) injections while they were pursued for 2 hours (10-12a.m) in the case of euthanasia. The amount of wakefulness, non-rapid eye movement (NREM)/ slow waves sleep (SWS) and REM sleep were quantified in baseline recordings as well as for 2h after all drugs injections and open field exposure.

### TRAPing

4-hydroxytamoxifen (4-OHT) was prepared as described previously (Lee et al., 2020). Briefly, 4-OHT (Cat# H6278 Sigma Aldrich, St. Luis, MO) was dissolved at 20mg/mL in absolute ethanol by ultrasonic water bath at 37°C for 10 min and was then aliquoted and stored at –20°C as a stock solution. Before injection, corn oil (Sigma Aldrich) was added to the thawed stock solution to replace the ethanol and to obtain 10 mg/mL 4-OHT, and then ethanol was evaporated at 37°C. The 10mg/mL 4-OHT solution was used the same day or one day after preparation. All animals were injected intraperitoneally (i.p.) with 80 mg/kg 4-OHT, 2h after the beginning of the first Wakefulness (W) induction.

### Wakefulness

Solriamfetol was dissolved in saline solution (NaCl 0.9%). Modafinil was dissolved in a vehicle containing dimethyl sulfoxide (DMSO) 3 % and sterile saline (NaCl 0.9 %) and was then injected immediately to each mouse. Drug solution was freshly prepared the day of the injection. Solriamfetol and modafinil were administered by intraperitoneal injection at a concentration of 10 ml/kg body weight at 10a.m. Wake onset was defined as the time elapsed between the time of injection and the first wake episode lasting at least 1 min and not interrupted by more than two 4 s epochs scored as NREM sleep. We did use 60 mg/kg for Solriamfetol (Sol) and 32 mg/kg for Modafinil (Mod) to obtain an induction of wakefulness (W) lasting around two hours. By comparison, Hasan et al. (Hasan et al., 2009) used 150 mg/kg for both Modafinil and Solriamfetol which are high doses possibly inducing non-specific side effects and jeopardizing conclusions regarding cFos activation. The NW protocol (open field) was used to induce wakefulness during 2h (before perfusion) or 4 hours for 4-OHT. In this case, 4-OHT was injected after two hours of wake and the mice were left in the open field two more hours to avoid TRAPing of neurons activated during sleep before the washout of the drug. When the mice were perfused, they were left in the open field for two hours before perfusion. To maintain the animals awake we put them in a round open field barrel containing wood tips and small objects. Food and water were freely available in the barrels. During exposure to the open field, the animals were permanently monitored by a web-camera from a different room to check whether they were staying awake. The animals were gently touched by a soft tissue when they became inactive/drowsy. Four groups of mice were generated. In the first group, solriamfetol was injected at 10a.m and 4-OHT was injected two hours after the beginning of the induction of wake. One week later, modafinil was injected and the mice were perfused after two hours of wakefulness (Sol-Mod group). In the second group of mice, the two protocols of waking induction were inverted (modafinil first and solriamfetol second, Mod-Sol group). In the third group, the animals were placed in the open field and 4-OHT was injected two hours after the beginning of the induction of wakefulness. One week later, solriamfetol was injected at 10a.m and the mice was perfused after two hours of wakefulness (NW-Sol group). In the fourth group of mice, the two protocols of wake induction were inverted (solriamfetol first and open field second, Sol-NW group). The mice were perfused 2h after being placed in the open field.

### Perfusion

All animals were deeply anesthetized with intraperitoneal (IP) injection of pentobarbital (140mg/kg) and were perfused with a heparin-added Ringer’s lactate solution (1:1000) followed by 4% paraformaldehyde/Phosphate Buffered Saline (PBS) (pH 7.4) for fixation. The brains were then post-fixed with 4% paraformaldehyde for one night at 4℃ and then were stored in 30% sucrose/PB for two days at 4℃.

### Immunohistochemistry

Brains were frozen with methylbutane placed on dry ice at around -30°C. Then the brains were sliced in 30 µm thick coronal sections serially distributed in eight wells at -20°C in a cryoprotective solution containing 20% glycerol and 30% Ethylene glycol in 0.05 M PB (pH 7.4). Brain sections were first washed in 0.1 M PBS with 0.4% Triton X-100 (Sigma-Aldrich) to remove the cryoprotectant. Sections were then incubated in 0.3% H2O2 for 1 hour to quench endogenous peroxidase activity, then washed 3*10 minutes in 0.1 M PBS with 0.4% Triton X-100. Different wells of brain sections were used to perform the different immunofluorescence protocols. They were all incubated with anti-c-Fos rat antibodies (1:50000, c-Fos antibody -226 017; Synaptic System), and incubated with either anti-Goat-Orexin (1:5000, Goat monoclonal IgG, sc-80263 Santa Cruz) or anti-Rabbit-TH (1:40000, Rabbit polyclonal, 213 102 Synaptic System) for 48 h at 4°C in 0.1 M PBS containing 0.4% Triton X-100. After being washed 3*10 minutes in 0.1 M PBS with 0.4% Triton X-100, sections were then incubated for 3 hours at room temperature in 0.1 M PBS with 0.4% Triton X-100 containing biotinylated rabbit Anti-Rat IgG antibody diluted to 1:1000 (Vector Laboratories), and donkey anti-goat IgG 647 nm antibody diluted to 1:500 (Invitrogen, A21447) for Orexin, or donkey anti-rabbit IgG 647 nm antibody diluted to 1:500 (Invitrogen, A31573) for TH. Then, they were washed 3*10 minutes with 0.1 M PBS with 0.4% Triton X-100. Following an incubation of 1h30 with streptavidin (SA)-HRP (Alexa Fluor™ Tyramide SuperBoost™ Kit, streptavidin; Life Technologies, 1:1000) in 0.1 M PBS with 0.4% Triton X-100, sections were washed 3*10 minutes in 0.1 M PBS with 0.4% Triton X-100, and incubated for 10 min in Alexa Fluor 488-conjugated Tyramide (Molecular Probes, Eugene, OR, USA) by diluting the stock solution 1:500 in 0.0015% H2O2/amplification buffer. The reaction was terminated after 10 min by rinsing the tissue in 0.1 M PBS. Following 3 washes with Phosphate Buffered Saline Tween (PBST) buffer, sections were mounted, dried, and cover-slipped with prolong Gold anti-fading reagent containing 4’,6-diamidino-2-phenylindole (DAPI) (Molecular Probes, Eugene, OR) and stored at 4 ℃. Sections were then imaged using an Axioscan Z.1 slide scanning microscope (Zeiss, Germany). Images were collected with a x20 objective (N.A. Plan-Apochromat 20×/0.8 M27 Air/0.8.) and a 0.45 Orca Flash camera. Dapi, c-Fos (Alexa-488), tdTomato (tdT), TH or hypocretin/orexin (Orx) (Alexa-647) were acquired using appropriate filter cubes, according to manufacturer recommendations. Exposure time was defined by signal distribution across gray values (>20% of measure gray values on 16 bits images). For all sections, five mosaics were acquired (5µm steps) then projected on a single plane, using the depth of focus “wavelet” function of the Zen software (version 3.1). Images were not modified and were directly imported in Neuroinfo (MBF Bioscience, USA) for quantification.

### Cell counting

The Allen Brain Reference Atlas (Adult Mouse) was used as reference for all structures. In triple-stained fluorescent sections, drawings of structures and automatic plotting of cFos+, tdTomato+ and Tyrosine Hydroxylase (TH) or Orexin/hypocretin (Orx) neurons was then made using a computerized image analysis system (Neuroinfo software, MBF Bioscience, USA). The cFos+, tdTomato+, double (cFos+/tdTomato+) and triple labeled neurons were automatically plotted and counted with the Neuroinfo software in all mice on 10 sections per mice with approximately 1mm spacing from the forebrain to the medulla oblongata. Cells were considered labeled when they exhibited clear cytoplasmic (for tdTomato, TH or Orx) or nuclear (for cFos) staining. Four representative mice per group (Sol-Mod; n=4, Mod-Sol; n=4, NW-Sol; n=4, Sol-NW; n=4) were analyzed. There was no statistical difference between hemispheres (not shown) and, therefore, counting from the left and right hemispheres were summed.

### Statistical analysis

Statistical analyses were conducted using Matlab (Mathworks, USA). Non parametric tests (Kruskal-Wallis followed by Mann-Whitney) were used to compare sleep-wake quantities after drug administration or NW protocol. Pearson’s correlation was employed to assess the association between the expression levels of tdT and cFos in individual animals. Data is organized by the following macroregions: cortex, telencephalon, thalamus, hypothalamus, hippocampal formation, mesencephalon, pons and medulla. A two-way analysis of variance (ANOVA) was applied to compare cell densities across different brain structures, considering two factors: the experimental condition (factor 1: Sol, Mod, and NW) and the staining type (factor 2: tdT and cFos), followed by the Tukey-Kramer post-hoc test. The Tukey-Kramer test is an extension of the Tukey Honestly Significant Difference (HSD) test that adjusts for unequal sample sizes while controlling the family-wise error rate (FWER) to prevent Type I errors during multiple pairwise comparisons. A similar approach was used to compare the percentage of TH neurons double labeled with cFos or tdT in all noradrenergic groups (NA). Paired t Tests were also performed to compare the density of tdT and cFos neurons between the first and second conditions in the same animals. Then, taking advantage of the TRAP2 model, the level of neuronal reactivation between the first and second experimental conditions was calculated. The Reactivation Index (RI) was obtained by dividing the number of tdT/cFos double-labeled neurons by the total number of tdT neurons for each structure. If all positive tdT cells (stained in the first condition) were recruited in the second condition, the number of double stained cells would be the same as the number of tdT neurons, and the RI would be equal to 1. On the other hand, if no overlap occurred between the second and the first condition, the RI would be equal to 0. We fitted the distribution of RI values in each one of the macroregions using a gamma distribution function to determine a reactivation threshold. A one-way ANOVA was performed to compare the RI between experimental groups or across macroregions, followed by the Tukey-Kramer post-hoc test. Statistical significance was defined at a threshold of 5% (p<0.05). Data are presented as mean ± SEM or median and interquartile range.

## Abbreviations

4-OHT: 4-hydroxytamoxifen
5HT: serotonin
μ: mean
ACB: nucleus accumbens
Ach: acetylcholine
ANOVA: Analysis of variance
BNST: bed nucleus of the stria terminalis
CEA: central amygdalar nucleus
CGRP: calcitonin gene-related peptide
CI: confidence interval
CRH: corticotropin releasing hormone
CTX: cortex
DA: dopaminergic
DAPI: 4’,6-diamidino-2-phenylindole
DMSO: dimethyl sulfoxide
DRN: dorsal raphe nucleus
EEG: electroencephalogram
EMG: electromyogram
FWER: family-wise error rate
GABA: gamma-aminobutyric acid
Glu: glutamate
Hcrt: hypocretin
HDC: histidine decarboxylase
HF: hippocampal formation
Hist: Histamine
HSD: Tukey Honestly Significant Difference test
HYP: hypothalamus
IP: intraperitoneal
LC: locus coeruleus
LDT: laterodorsal tegmental nucleus
LHA: lateral hypothalamic area
LPB: lateral parabrachial nucleus
LRN: lateral reticular nucleus
LS: lateral septum
MCH: melanin-concentrating hormone
Md: median
MED: medulla
MES: mesencephalon
Mo: mode
Mod: Modafinil
MPO: medial preoptic nucleus
NA: noradrenergic
NDRI: norepinephrine–dopamine reuptake inhibitors
NTS: nucleus of the solitary tract
NREM: non rapid eye movement
NW: non-pharmacological wakefulness
Orx: orexin
OT: oxytocin
PBS: Phosphate Buffered Saline
PBST: Phosphate Buffered Saline Tween
PCA: Principal Component Analysis
PON: pons
PS: paradoxical sleep
PSTN: parasubthalamic nucleus
PVN: paraventricular hypothalamic nucleus
PVT: paraventricular thalamic nucleus
REM: Rapid Eye Movement
RI: Reactivation Index
SEM: Standard error of the mean
SN: substantia nigra
SON: supraoptic nucleus
SOI: structures of interest
Sol: Solriamfetol
SSp: somatosensory
SWS: slow waves sleep
TAAR1: trace amine associated receptor 1
tdT: tdTomato
TEL: telencephalon
TH: tyrosine hydroxylase
THA: thalamus
TMN: tuberomammillary nucleus
TRAP2: targeted recombination in active populations
VLPO: ventrolateral preoptic area
VP: vasopressin
VTA: ventral tegmental area
W: wakefulness
ZI: zona incerta

## Acknowledgements

This work is supported by CNRS (UMR5292),INSERM (U1028), SFRMS, University Lyon 1, and Coordination for the Improvement of Higher Education Personnel (CAPES). We also thank the CIQLE platform of the SFR Santé Lyon Est for scanning the sections. This work was performed within the framework of the LABEX CORTEX(ANR-11-LABX-0042) of l’Université Claude Bernard Lyon 1, within the program “Investissements d’Avenir” (decision n° 2019-ANR-LABX-02) operated by the French National Research Agency (ANR).

